# Spatial distribution of bacteria and extracellular polymeric substances impacts nanoparticle penetration in biofilms

**DOI:** 10.1101/2022.06.14.496116

**Authors:** Bart Coppens, Tom E. R. Belpaire, Jiří Pešek, Hans P. Steenackers, Herman Ramon, Bart Smeets

**Affiliations:** Division of Mechatronics, Biostatistics, and Sensors, KU Leuven, 3001 Leuven, Belgium; team SIMBIOTX, Inria Saclay, 91120 Palaiseau, France; Centre for Microbial and Plant Genetics, KU Leuven, 3001 Leuven, Belgium

**Keywords:** Biofilm architecture, Nanoparticle treatment, Brownian dynamics model, Single Particle Tracking

## Abstract

Extracellular polymeric substances (EPS) in bacterial biofilms complicate treatment by inactivating drugs and slowing down diffusion. Through enhanced penetration and resistance to degradation in bacterial biofilms, nanoparticle (NP) carriers can help improve biofilm treatment. However, the way in which biofilm architecture influences the diffusive properties and penetration of NPs in biofilms is still poorly understood. In this work, we combined single particle tracking (SPT) and confocal laser scanning microscopy (CLSM) in *Salmonella* biofilms with simulations of a Brownian dynamics model to quantify how macro- (spatial organization of the bacteria) and micro- (EPS dependent) structure of the biofilm affects NP penetration. In CLSM images we observed immobilization of NPs in the EPS, which allows shielding of bacteria from the NPs, an effect that was more pronounced in dispersed biofilms, grown in nutrient-rich conditions, than in compacted biofilms, grown in nutrient-poor conditions. SPT experiments revealed anomalous diffusion, with an increased probability for small displacements near clusters of bacteria. Simulations of a Brownian dynamics model revealed that EPS reinforces shielding by affecting the pore structure of the biofilm. Finally, in virtual biofilms with varying spatial distribution of bacteria, we found that even for the same number of bacteria, dispersed biofilm structures provide more shielding than biofilms organized in dense, compacted clusters, even when accounting for decreased NP diffusivity.

## Introduction

Biofilms are communities of bacteria, typically encapsulated in a self-produced matrix of extracellular polymeric substances and considered to be the predominant bacterial lifestyle (1, 2). The EPS form a barrier that hampers treatment with conventional antibiotics by slowing down diffusion or inactivating them (3, 4). A possible strategy to overcome these issues is to encapsulate antibiotics in liposomal or polymeric NPs (3–6). NPs act as carriers that protect antibiotics from inactivation and can be modified to enhance biofilm treatment. Changing particle size can alter penetration through the porous structure of the matrix, adding differently charged groups can affect electrostatic interactions with the mainly negatively charged matrix, and functionalization with specific chemical groups can improve specificity (5–8). As versatile drug-delivery systems, NPs have already shown great promise as antimicrobial strategies (3, 5) and are anticipated to play an important role in combating the surging problem of antibiotic tolerance in biofilms (9).

Delivery of NPs within biofilms, however, faces challenges inherent to diffusion in mucus-like environments, such as low permeability, high retention and formation of protein coronas (10, 11). NP movement through the biofilm matrix is influenced by three main types of interactions: (i) size-dependent filtering due to limited pore size (10, 12–14), (ii) electrostatic interaction from charged components in the biofilm matrix (10, 12–15), and (iii) chemical interactions (11, 16, 17). The effect of these interactions on NP diffusion is non-trivial. Binding interactions often hinder penetration, leading to phenomena such as subdiffusive behaviour, where diffusion is slowed down, such that their mean squared displacement (MSD) scales as ⟨*x*^2^(*t*)⟩ ∝*t*^*α*^, where 0 < *α* < 1 (10, 12, 18). However, the existence of weak interactions with the matrix can also enhance NP perfusion (16). A suitable technique to study these interactions and their effect on NP fate is single particle tracking (SPT). In SPT, fluorescent NPs probe the biofilm micro-environment non-invasively, so that local mechanical properties can be inferred from particle diffusion parameters (19–22), and the penetration capability of a variety of NPs can be assessed (12, 19, 21).

Interactions between the EPS matrix and NPs are therefore recognized as important recalcitrance mechanisms for the biofilm (4,5, 7, 8, 23). However, the impact of the spatial distribution of this EPS matrix, along with the spatial organisation of the biofilm bacteria on which the EPS is contingent, has not been extensively studied so far. Nonetheless, this so-called “biofilm architecture” directly influences (i) pore sizes, which get smaller close to dense groups of bacteria (24), (ii) connectivity within the biofilm (25) and (iii) spatial heterogeneity of diffusion coefficients (14). However, while biofilm architecture can be controlled partially in the lab *in vitro* (4, 26, 27), it is impracticable to fully separate structural effects from other biofilm properties such as matrix composition, since these are strongly linked (28). To this end, computational simulations may provide complementary understanding (7, 8). Continuum approaches have provided insight into diffusion mechanics, for example using a plumpudding model (29), or in general reaction-diffusion models (30). In crowded environments such as biofilms, particle-based Brownian dynamics models are valuable, as they allow for varying particle shape and surface properties, grant insight in processes impacting diffusion, give single-particle information and can be directly compared to experimental SPT and CLSM data (19, 31–35). Moreover, interactions between NPs and EPS can be modelled explicitly (31, 32, 36), or they can be coarse-grained and represented implicitly (37), allowing for computationally cheaper, large scale simulations. To allow these latter type of simulations, representative biofilm structures first need to be generated either via biofilm growth (38) or via continuum equations (39, 40).s

In this work, we investigated the impact of biofilm architecture by studying how (i) spatial distribution of bacterial clusters, or “macro-structure” and (ii) the amount of EPS produced by the bacteria within the clusters, or “micro-structure” affect the penetration of NPs in *Salmonella enterica* Typhimurium biofilms. By changing nutrient availability, we controlled the degree of compaction/dispersedness of *in vitro* biofilms in order to influence macro-structure. Using fluorescence microscopy we tracked and localized individual NPs in these biofilms to quantify their diffusion properties and penetration. Furthermore, to investigate the effect of micro-structure for different types of macro-structure, we performed simulations of a Brownian dynamics model of NP diffusion in biofilms. These simulations revealed that more dispersed biofilm structures show increased shielding of bacteria from diffusing NPs, relative to compact biofilm structures.

## Results

### Structurally distinct biofilms differ in nanoparticle penetration

We started by studying the impact of the biofilm macro-structure on nanoparticle penetration. In an attempt to establish different types of macro-structure, biofilms of *S*. Typhimurium were grown in high and low nutrient environments for 48 h and analyzed with CLSM. The abundance of nutrients appeared to have a pronounced effect on biofilm structure: In nutrient-poor conditions, we observed a lower global bacterial density (3.2 ± 1.0 v/v%), and thus associated biomass, compared to nutrient-rich conditions (8.3±2.9 v/v%), see Supplementary Table 1. Additionally, bacteria appeared more compacted in nutrient-poor conditions (Fig. 1a-b). We quantified the degree of compaction via the average pairwise distance of each bacterium to their 10 nearest neighbours. Nutrient-poor biofilms showed a higher compaction with an average pairwise distance of 2.50 ± 0.07 µm, while nutrient-rich biofilms showed an average pairwise distance of 2.75 ± 0.05 µ m. Such differences in macro-structure can be expected to impact NP penetration. On the one hand, increased biofilm compaction will increase the volume of pores and might thereby enhance NP penetration through the biofilm. On the other hand, more dense, compact clusters might inhibit NP from entering and thereby protect bacteria at the center of the cluster. We therefore used CLSM to determine the three-dimensional position of confined aminated and carboxylated fluorescent NPs (10^−5^ w/v%) 1 h after introduction in 48 h old biofilms. NPs were found to be more concentrated in the upper layers of the nutrient-rich grown biofilms and unable to penetrate to the bacteria close to the substrate, while for nutrient-poor biofilms they were present closer to the bacterial clusters. To quantify NP penetration in the biofilm, we introduced the “affinity” and “coverage length” measures, Fig. 1c. High affinity, which we defined as the percentage of NPs closer than a threshold of 0.3 µm from the bacterial surface, indicates the ability of NPs to reach bacteria in the biofilm. Dependency of affinity on this threshold is shown in Supplementary Fig. 1. The value of this threshold was informed by matrix staining Fig. 1d, where the matrices in nutrient-poor biofilms reach on average 0.31 ± 0.07 µm from the surface of the bacteria and 0.27 ± 0.05 µm for nutrient-rich biofilms. In treatment, high affinity results in more specific drug release and a lower required dose of NPs, Fig. 1e. We found no significant differences in affinity between biofilms grown in rich and poor nutrient conditions, both for aminated and carboxylated NPs. Affinity was, however, significantly lower for aminated than for carboxylated NPs in nutrientrich biofilms (*p* = 0.006). As a second measure, we defined the coverage length as the median distance from each bacteria to the closest NP, Fig. 1e (full distributions are shown in Supplementary Fig. 2). We found a 5-fold higher coverage length in nutrient-rich grown biofilms compared to nutrient-poor grown biofilms, for both aminated (*p* = 0.004) and carboxylated (*p* = 0.002) nanoparticles. One evident factor that partially explains this difference in coverage length is the discrepancy in total biomass, since nutrient-rich conditions had approximately 2.3 fold higher biomass compared to nutrient-poor conditions. Moreover, it is possible that reaction-diffusion mechanisms changed the chemical micro-environment in the biofilm depending on nutrient-availability, leading to altered gene expressions and thus possibly different EPS matrix properties (23, 30). In order to understand the impact of biofilm architecture on NP penetration, independent of the confounding effects of biomass, it is thus valuable to first quantify the diffusion barriers that hinder NP movement at the micro-structural scale.

**Fig 1.**
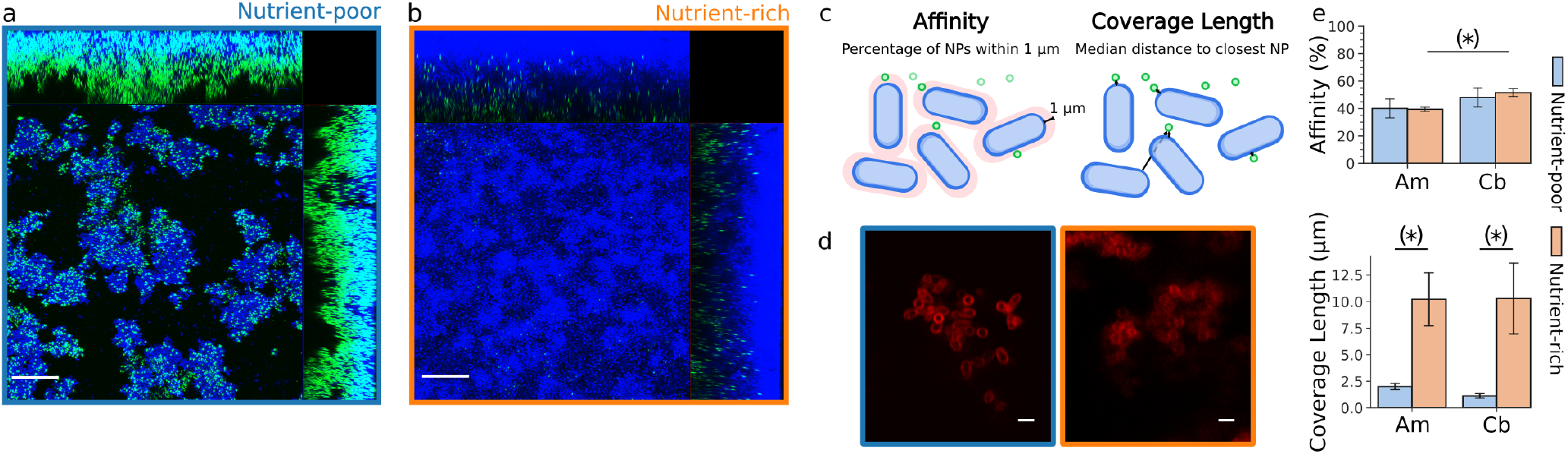
Overview of the 3D microscopy of nanoparticles incubated in a *Salmonella enterica* Typhimurium biofilm. Nanoparticles are green fluorescently labeled, either aminated (Am) or carboxylated (Cb), with blue fluorescently labeled bacteria, either cultivated in nutrient-rich or nutrient-poor conditions. **a** Max intensity orthogonal projection of a nutrient-poor biofilm with aminated nanoparticles. **b** Max intensity orthogonal projection of a nutrient-rich biofilm with aminated nanoparticles. **c** Schematic representation of affinity and coverage length measures. **d** CLSM images of red fluorescently stained curly and cellulose for nutrient-poor and nutrient-rich biofilms. Data shown for different biofilms than a, b and e. **e** Results for affinity and coverage length. Error bars are the standard deviation of three biological repeats. Significance levels are tested with pairwise Student’s t-tests, without multiple correction. (top) (∗) is a p-value of 0.006, (bottom) (∗) are p-values of 0.004 for Am and 0.002 for Cb.

### Nanoparticle diffusion in *Salmonella* biofilms is heterogeneous

To elucidate the processes that govern NP penetration, we performed SPT to quantify the diffusion characteristics of fluorescent NPs within the nutrient-poor 48 h old *Salmonella* biofilms. We found a range of different dynamic behaviors of NPs depending on their proximity to biofilm bacteria and the architectural elements the NPs are exposed to. Whereas NPs in the pores between clusters of bacteria move more freely, NPs near the clusters appear more confined and move more slowly, Fig.2a. We observed a similar disparity based on the ensemble displacement distribution, where there is an increased probability of small displacements, indicative of confined movement, Fig.2b. The empirical ensemble displacement distribution (DispD), with full data in Supplementary Fig. 3, shows exponential tails, further deviating from a Gaussian distribution that would be characteristic of simple diffusion. In case the diffusion coefficient *D* follows an exponential distribution, the ensemble distribution of displacements is expected to exhibit such exponential tails. This effect can, however, even occur for a non-exponential distribution of *D* at sufficiently short lag times (22, 41). In our results, the measured effective diffusion coefficient, calculated from the time-averaged MSD, instead follows a bimodal distribution, with a relatively small fraction of mobile particles and a large fraction of immobilized particles for which D ≈ 0, Fig. 2c. Finally, the diffusion exponent *α*, assuming 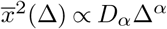, indicates the presence of subdiffusive anomalous diffusion Fig. 2d, where *α* < 1 and NPs are confined to a small area. Overall, the observation of strongly heterogeneous mobility substantiates the impact of macro-structural elements on nanoparticle penetration. In addition, the low mobility of a large fraction of NPs emphasizes that confinement in the EPS micro-structure is an important factor in hampering NP penetration.

**Fig 2.**
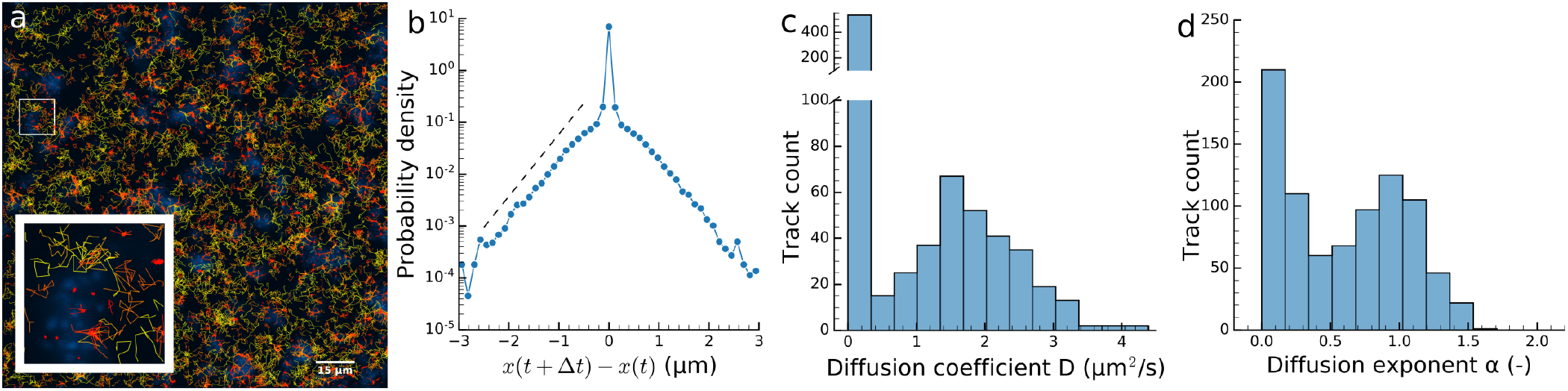
Overview of SPT tracks and data. Data shown for aminated polystyrene nanoparticles after 20 minutes incubation in *Salmonella enterica* Typhimurium biofilm. **a** Visualization of the analyzed tracks. Bacteria are presented in blue and were imaged separately before the tracks were imaged. Nanoparticle tracks are coloured according to their mobility. **b** Ensemble probability distribution of displacements with 0.1 s lag time. The dashed line shows the slope for a Laplacian fit, estimated via non-linear least squares on the displacement curves on log-scale. **c** Distribution of the diffusion coefficients *D*, estimated via linear least squares on the time averaged mean squared displacement (TAMSD) as 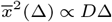.A break was included in the y-axis to show the distribution of larger *D* as well. **d** Distribution of anomalous diffusion exponents *α*, estimated via nonlinear least squares on the TAMSD.

### EPS thickness differentially impairs nanoparticle penetration by tuning percolation

To further study the effects of macro-structure and their interplay with micro-structure, we simulated the diffusion of NPs in a biofilm with a particle-based Brownian dynamics model. This model allows to consider bacteria organized according to different macro-structures informed by microscopic images, with different amounts of EPS to manipulate the micro-structure. Bacteria are represented via spherocylinders, whereas NPs are represented as spheres. NPs only interact with bacteria, not with each other, and we assumed bacteria are static with respect to NPs. EPS is represented via a Gaussian-shaped viscosity kernel *η*(***x***_*i*_), with a high viscosity Δ*η*_*M*_ near the surface of bacteria, declining to bulk viscosity *η*_0_ as

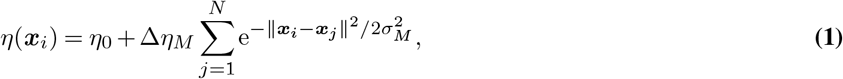

with ***x***_*j*_ the position of bacteria *j*. Motivated by the observation that NPs are immobilized strongly inside dense clusters of bacteria, we considered the viscosity kernels as additive in our model, Fig. 3a. This assumption is consistent with diffusion analyses in biofilms, which revealed a lower diffusivity in larger cell clusters, further decreasing in denser parts of clusters (14, 24). Our coarse-grained strategy to model NP-EPS interactions permits simulation of NP penetration in large biofilm systems in a computationally favorable manner. Varying the EPS viscosity Δ*η*_*M*_ and the length-scale of the EPS *σ*_*M*_ allows us to tune the diffusion characteristics of the EPS structure in simulations. We estimated the effective thickness of EPS in experimental conditions as ≈ 0.3 µm, Fig. 1d, thereby informing the range of variation in the parameter study of *σ*_*M*_. Next, we simulated the diffusion of NPs in the biofilm structures obtained from image segmentation of both compacted, nutrient-poor biofilms and dispersed, nutrient-rich biofilms, Fig. 1. Since we did not observe a significant difference in penetration between aminated and carboxylated NPs, we did not explicitly differentiate between the two in our simulations. Examples of segmented biofilms with simulated NPs are shown in Fig. 3b-c. For these structures, increasing the EPS characteristic length *σ*_*M*_ results in a decrease in pore volume where NPs can diffuse freely. Figs. 3b-c, bottom left, show the decline in pore volume for a nutrient-poor biofilm when *σ*_*M*_ increases from 0.5 µm to 2.0 µm. In contrast to nutrient-rich biofilms, Fig. 3b, the nutrient-poor biofilm remains a percolated system. The DispD for the nutrient-poor biofilm, Fig. 3b-c, bottom right, shows similar profiles to the profile obtained from SPT experiments, Fig. 2b. As EPS viscosity Δ*η*_*M*_ increases, a transition occurs from a Gaussian DispD to a non-Gaussian DispD with increased probability for short displacements, Fig. 3d, The width of the central peak in this non-Gaussian distribution increases and the probability of large displacements decreases with *σ*_*M*_, Fig. 3b, bottom right, an effect that is even more pronounced for the nutrient-rich biofilm, Fig. 3c, bottom right. Indeed, the connectivity of the pore structure drops with increasing *σ*_*M*_ as percolation vanishes, and entrapment of NPs is observed, resulting in strongly non-Gaussian tails in the DispD. The affinity measure in Fig. 3e was calculated with the same threshold 0.3 µm as in Fig. 1e. Affinity of NPs to bacteria was found to first increase with characteristic EPS length-scale *σ*_*M*_ as the probability to interact with the biofilm increases, and then drop for *σ*_*M*_ > 0.2 µm, when the matrix increasingly separates the NPs from the surface of the bacteria. Affinity was lower in nutrient-poor conditions than in nutrient-rich conditions, but only for small values of *σ*_*M*_. Considering coverage length (full distributions available in Supplementary Fig. 4), a first observation is that increasing Δ*η*_*M*_ from Δ*η*_*M*_ = 1 mPas onward leads to a small decrease in coverage length, Fig. 3f, since NPs are now immobilized closer to bacteria, without inhibiting their penetration altogether as they can still escape. This confirms previous studies that found that some weak interactions might benefit penetration (16). Higher viscosities, however increase the coverage length due to stronger NP immobilization. The coverage length increases with *σ*_*M*_, which is more pronounced for nutrient-rich than for nutrient-poor biofilms, Fig. 3g and, similar to our experiments, the coverage length in nutrient-rich biofilms is higher relative to nutrient-poor biofilms. The effect where *σ*_*M*_ increases coverage length more in nutrient-rich biofilms cannot be attributed merely to pore volume, as nutrient-rich biofilms with lower *σ*_*M*_ have similar pore volume fraction as nutrient-poor biofilms with higher *σ*_*M*_, see Supplementary Fig. 5 and Fig. 6, while the coverage length remains higher for nutrient-rich biofilms. Due to decreased percolation in the pore structure of nutrient-rich biofilms, Fig. 3b, NPs may become trapped in the upper layers of the biofilm. This comparison of coverage length at similar effective pore volume thus further supports that the spatial distribution of cells, i.e. macro-structure, in interaction with the local EPS, plays an important role in determining NP penetration within the biofilm, Fig. 3b, c, f and Supplementary Fig. 5.

**Fig 3.**
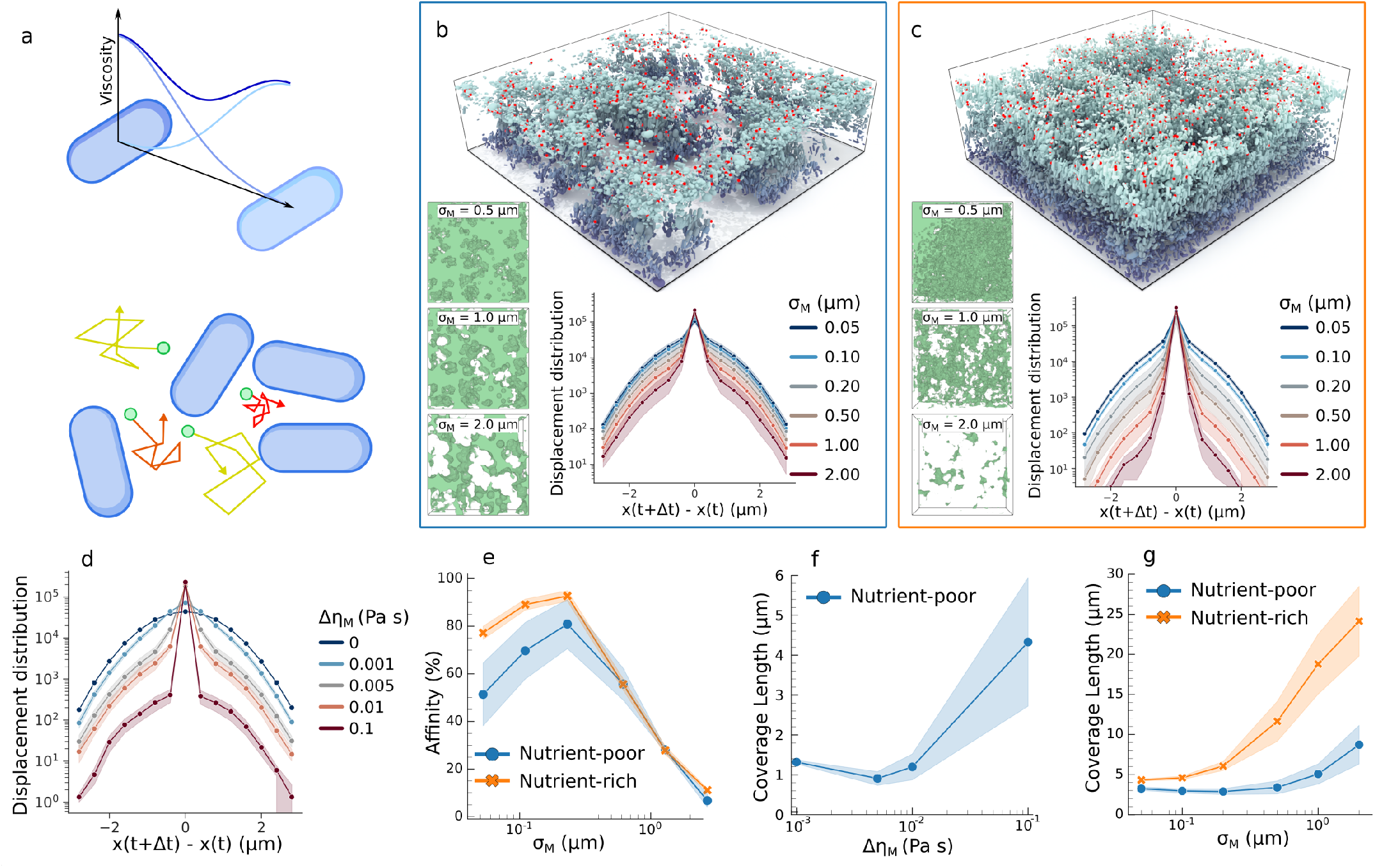
Simulations within segmented *in vitro Salmonella enterica* Typhimurium biofilms. **a** Schematic representation of the Gaussian viscosity kernel in our Brownian dynamics model. Nanoparticle (green) tracks are coloured relative to viscosity, which depends on proximity to bacteria (blue). **b** Simulated nanoparticles (red) in segmented nutrient-poor biofilm (133 × 133 × 34 μm), coloured relative total biofilm height. (left) XY projection of channels where *η* = *η*_0_ in a nutrient-poor biofilm. (right) Ensemble displacement distribution of simulated nanoparticles after diffusing for 10 minutes. **c** Simulated nanoparticles (red) in segmented nutrient-rich biofilm (133 × 133 × 34 μm), coloured relative total biofilm height. (left) XY projection of channels where *η* = *η*_0_ in a nutrient-rich biofilm. (right) Ensemble displacement distribution of simulated nanoparticles after diffusing for 10 minutes. **d** Ensemble displacement distribution of simulated nanoparticles after diffusing in a segmented nutrient-poor biofilm for 10 minutes, coloured according to Δ*η*_*M*_. **e** Affinity for simulated nanoparticles, closer than 1 μm to segmented bacteria in nutrient-poor and rich biofilms, for varying *σ*_*M*_, at constant Δ*η*_*M*_= 10 mPa s. Filled area is the standard deviation in three segmented biofilm biological repeats. **f** Coverage length for simulated nanoparticles in nutrient-poor biofilms for varying Δ*η*_*M*_, at constant *σ*_*M*_= 0.5 μm. Filled area is the standard deviation in three segmented biofilm biological repeats. **g** Coverage length for simulated nanoparticles in nutrient-poor and rich biofilms for varying *σ*_*M*_, at constant Δ*η*_*M*_= 10 mPa s. Filled area is the standard deviation in three segmented biofilm biological repeats.

### Dispersed biofilm architectures provide shielding that limits nanoparticle penetration

Although our previous analyses clearly hint at an impact of biofilm architecture on NP penetration, they were all confounded by differences in total number of cells. To explicitly assess how spatial macro-structure of the biofilm, apart from its total biomass, affects the penetration success of NPs, we apply our model to virtual biofilms that differ in spatial cell distribution but conserve the total number of bacteria. To this end, we solved the Cahn-Hilliard equations (CH) for phase separation for a binary mixture (void space and biofilm). Phase separation theory has been used to characterize experimental biofilm formation (39, 40), and consequently applied to generate representative virtual biofilm structures (42). As the mixture coarsens over time *t*, the characteristic length of domains increases as *L*_*t*_ ∼ *t*^1/3^. Using a zero concentration boundary condition at the top and natural boundary conditions at the remaining sides, we obtained a collection of virtual biofilms with varying degrees of compaction but with an equal number of bacteria, Fig. 4a. These structures vary from a near uniform distribution of bacteria at low *L*_*t*_ to highly compacted biofilm structures at high *L*_*t*_. The pore structure for *σ*_*M*_ = 0.4 µm and *σ*_*M*_ = 1.0 µm is shown in Fig. 4b. At high *L*_*t*_, the pores between clusters are large, and pore structure remains percolated even at large *σ*_*M*_. However, the viscosity within compact clusters at high *L*_*t*_ is greatly elevated due to the additive nature of the viscosity kernel, see Supplementary Fig. 7. Simulations of NP diffusion in these structures show that affinity is reduced for increasing biofilm compaction, as more NPs diffuse freely in the larger open pore space, Fig. 4c. We found that coverage length decreases with *L*_*t*_, Fig. 4d, an effect that is more pronounced at large *σ*_*M*_, see Supplementary Table 2 and Supplementary Fig. 8. For small *L*_*t*_, the network of channels reaches the percolation threshold for increasing *σ*_*M*_, while for large *L*_*t*_, the system remains percolated. Moreover, these results highlight the capacity of more spatially distributed and loose biofilm structures to act as a “sieve” to retain diffusing particles. Their greater surface-to-volume ratio permits efficient absorption of particles, thereby impeding them from penetrating more deeply in the biofilm structure. In case of antimicrobial treatment using NPs, the NP acts as a source from which antibiotics can diffusive outward. To demonstrate the difference in antibiotics release from a NP in thick and compact versus thin and sparse biofilm structures, we simulated diffusion from a point-source — the hypothetical NP — in a gyroid solid of different length scale (*L*_*g*_), taking into account a decrease in diffusivity and a fixed absorption rate in the solid phase, Fig 4e. These simulations show that the main effect of thinner, more dispersed structures is a more concentrated dose near the source of the antibiotics, which is more diluted for larger *L*_*g*_. However, further from the source, the difference in the distribution of the relative dose vanishes.

**Fig 4.**
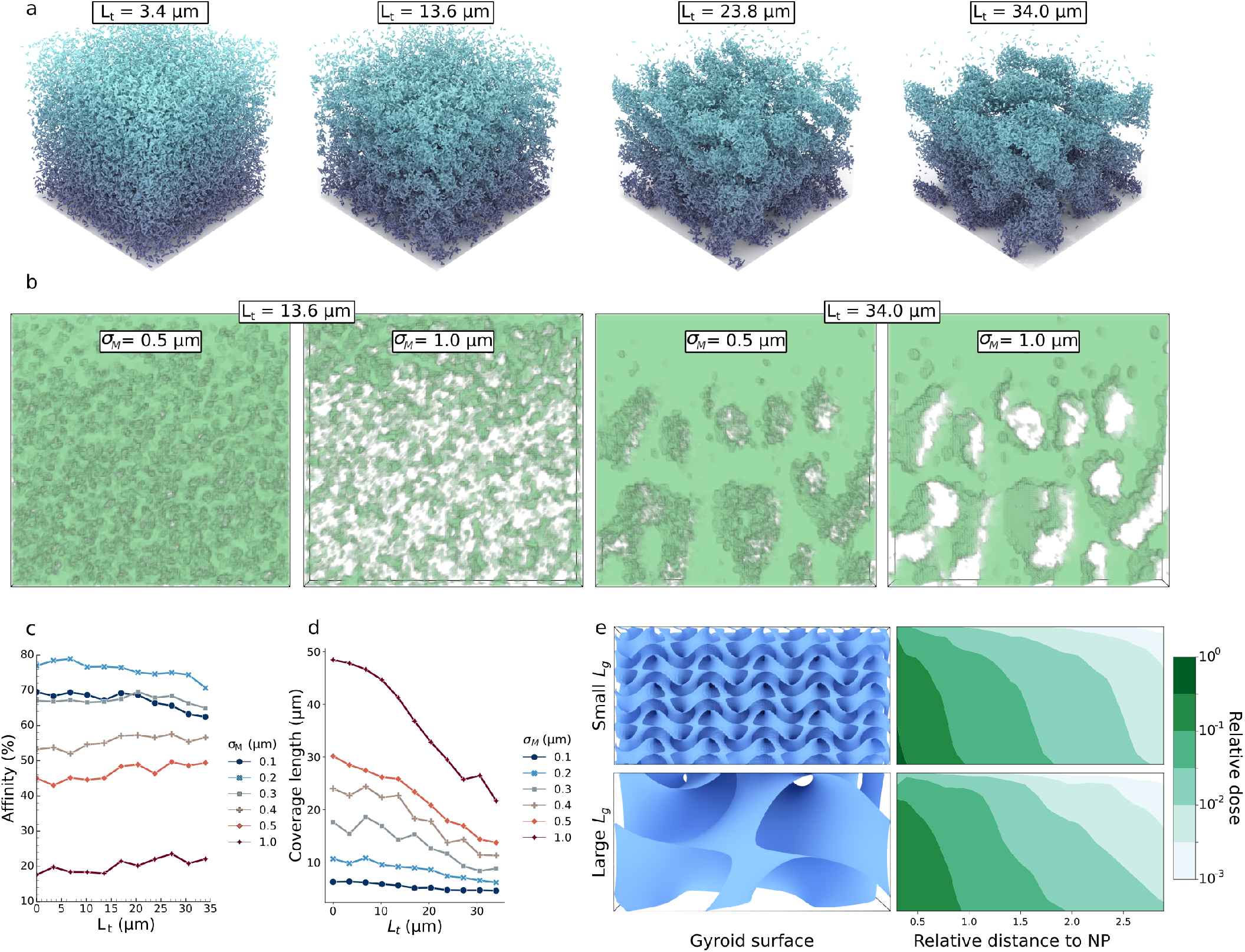
Results for simulated virtual biofilms, generated from Cahn-Hilliard equations for phase separation. **a** Virtual biofilms for various characteristic lengths *L*_*t*_. The biofilms have dimensions of 133 × 133 × 133 μm. **b** XZ views of channels in our virtual biofilms. For visual purposes we visualized only a slice of 20 μm, at the center of the biofilm. **c** Affinity for nanoparticles closer than 1 μm in virtual biofilms, at constant Δ*η*_*M*_= 10 mPa s. **d** Coverage length for nanoparticles in virtual biofilms, at constant Δ*η*_*M*_= 10 mPa s. **e** (left) Gyroid surfaces used in our FEM simulations. At the surface of these gyroid surfaces, absorption is high and diffusivity is low. (right) Absorbed concentration of antibiotics relative to their source concentration for high and low gyroid characteristic length *L*_*g*_, shown as distributions.

## Discussion

Although encapsulation of antibiotics in NPs protects the compounds from immobilization and inactivation (3), interactions between NPs and the EPS can still lead to poor penetration in the biofilm (7, 8, 16). While the nature of these interactions has been extensively studied (4, 7, 8, 16), the effect of biofilm architecture on NP penetration capabilities has received little consideration.

With three-dimensional CLSM, we located carboxylated and aminated nanoparticles in nutrient-rich and nutrient-poor biofilms. Although a higher affinity was expected for aminated NPs due to the presence of negatively charged elements in the biofilm (4, 7, 43), we did not observe significant differences in penetration capabilities between aminated and carboxylated NPs. In previous studies, positively charged NPs showed higher binding capabilities, lower diffusion coefficients and were generally more restricted in penetrating through the biofilm (12, 44, 45). We hypothesise that in our experiments, the size of the NPs contributed more to their capture than their surface charge, which would explain differences between our results and experiments using charged quantum dots (17). Moreover, since we measured NP position after one hour of incubation, differences in penetration due to differences in electrostatic interactions might also have aged at that moment. We observed that nutrient-rich biofilms exhibited a decrease in penetration capabilities, which can be attributed to the decrease in compaction and consequent increase in volume, occupied by bacteria and EPS (SI Table 2), consistent with plumpudding models (29) and other reaction-diffusion models (30). The observed increase of this biovolume in nutrient-rich biofilms implies more binding sites, bacterial clusters or ‘plums’, which can capture the diffusing NPs in the plumpudding, thereby preventing further penetration (29).

In SPT experiments of NPs in *Salmonella* biofilms, we found that a large number of NPs got immobilized or confined near clusters of bacteria, where the matrix is denser and less porous (14, 24). Moreover, we observed anomalous diffusion, characterized by exponential tails in the displacement distribution, which might be attributed to an underlying distribution of diffusion coefficients of non-immobilized NPs (22, 41). Consequently, NPs cannot be simply categorized as either immobile when close to clusters of bacteria or freely moving when they diffuse further from bacteria. Instead, NPs moving further from bacteria experience a variety of steric, electrostatic or chemical interactions, leading to highly heterogeneous diffusion coefficients. In *Salmonella* biofilms, we found diffusion coefficients for NPs with 60 nm radius, ranging between 0 and 4 µm^2^*/*s, which is compatible with other values reported in literature (12, 18, 19). The observed subdiffusion is not necessarily detrimental for biofilm treatment. NPs are often designed to have high affinity for bacteria, such that release of their contents happens close to the target. Yet, biofilm eradication is hampered when interactions with the matrix inhibit NPs from reaching groups of bacteria, although weak interaction might even enhance penetration and drug-delivery capabilities (16).

In order to analyze the effect of matrix transport properties in these structures in isolation, we simulated NP diffusion using a Brownian dynamics model. This model was able to represent the characteristic signature of heterogeneous diffusion observed in the SPT experiment. As such, it can provide a powerful alternative to continuum-type models for simulating diffusion in strongly heterogeneous or anomalous environments, while also furnishing information on individual particle trajectories and anomalous diffusion characteristics. Simulations in segmented biofilms predicted that NP affinity to bacteria first increases with increasing EPS thickness until it drops at very large thickness. From a treatment perspective, this provides an interesting trade-off, as it shows that the presence of matrix possibly benefits drug delivery by immobilizing NPs closer to bacteria. On the flip side, we found that increasing EPS thickness mainly has an adverse effect on the coverage length, by shielding more bacteria from NPs. This effect is particularly strong in high nutrient conditions, when the pore space reaches the percolation threshold at higher cell density.

Finally, we studied the effects of micro- and macro-structure separately, without confounding effects of biomass, in virtual parameterized biofilms generated using the Cahn-Hilliard equations. These simulations show that, at equal biomass, bacteria are better shielded from NPs in disperse biofilms compared to dense compacted biofilm structures, even when taking into account a proportional decrease in diffusivity in clusters that are more compacted. This suggests that the appropriate conceptual model to understand NP penetration is the model of a particle ‘sieve’ or a ‘filter’. Through its heterogeneous diffusion environment, the EPS provides an absorbing surface that effectively filters NPs, preventing them from penetrating further in the biofilm. From the evolutionary perspective of bacteria, the colony is better protected against chemical stress by growing sparsely and vertically, as long as the affinity between the chemical stressor and EPS is high. In these conditions, protection is provided through a large surface-to-volume ratio at the sacrificial upper layers rather than by a large size of individual cell clusters.

In the context of delivery of antibiotics through NPs, these results can be further clarified by considering the Thiele modulus 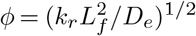 (46), where *k*_*r*_ is the sorption rate coefficient of the antibiotic, *D*_*e*_ its effective diffusion coefficient and *L*_*f*_ the characteristic diffusion length, which we can interpret as the coverage length if the source of antibiotics is the NP. When *ϕ* is small, diffusion is fast compared to reaction and the full material is treated. Conversely, when *ϕ* is large, reaction dominates and treatment is expected to be uneven. Hence, one expects large values of *L*_*f*_ for disperse biofilms and small values of *L*_*f*_ in compact biofilms due to better NP penetration. However, *D*_*e*_ is also likely to be higher in disperse biofilms, since the typical path between an absorbed NP and bacteria is more porous. Yet, the dominant parameter is expected to be the length-scale *L*_*f*_, as physical separation trumps diffusion barriers. The latter idea is further supported by the results of reaction-diffusion simulations from a point-source inside gyroid structures of varying coarseness, which show that apart from more dilution in local dose near the point source, the presence of larger structures has little effect on the spatial distribution of relative dose.

Our work has some limitations that warrant discussion. First, we consider the diffusion of NPs in the absence of convection by external fluid flows. However, fluid flow is expected to be an important factor in both shaping biofilm architecture and influencing nanoparticle transport (38, 47–49). Effects of convective flows and change in percolation structure due to bacterial motility are also disregarded, as we consider biofilms to be fully static at the time-scale of NP diffusion. Second, NPs are considered to be non-interacting in the simulations. Aggregation as a result of Van der Waals or electrostatic interactions could instigate additional size effects due to accumulation, further complicating NP penetration (8, 50). Third, the seemingly exponential tails of the experimental DispD do not emerge from our simulations. Addition of a long-range Gaussian kernel, could result in exponential-like tails of the DispD of the simulations. Finally, antibiotics can heavily impact biofilm architecture through alteration of bacteria-bacteria interactions and localized killing of bacteria (51, 52). Insights in the continual feedback between biofilm architecture and NP fate are crucial for the future application of NP-based antimicrobial delivery strategies and thus confer interesting research perspectives. Even so, we were able to show the importance of biofilm architecture on NP fate, demonstrating that even for constant bacterial density, their spatial ordering contributes greatly to shielding bacteria away from NPs.

## Materials & Methods

### Bacterial strain and growth conditions

The constitutive promoter PLλ and the fluorescent protein mtagBFP2 (53) were cloned into the multiple cloning site of the pFPV25 plasmid, kindly provided by Raphael H. Valdivia and Stanley Falkow (54), via restriction digestion. All primers used for the construction of this plasmid are listed in Table 1. Restriction enzymes were purchased from Roche and used according to the instructions of the manufacturer. *Escherichia coli* DH5α and *Escherichia coli* Top10F’ were used for cloning steps. The new construct were verified by sequencing and subsequently electroporated into *Salmonella entirica*, subsp. *enterica* serovar Typhimurium ATCC14028 using a Bio-Rad gene pulser.

Overnight cultures (ONC) were grown at 37 °C in Lysogeny broth (LB) in test tubes while shaking at 200 RPM. For cloning, colonies were grown on LB plates containing 1.5% agar (w/v). If the pFPV25 was present, 100 µg/mL was added both ONC and plate cultures.

**Table 1.**
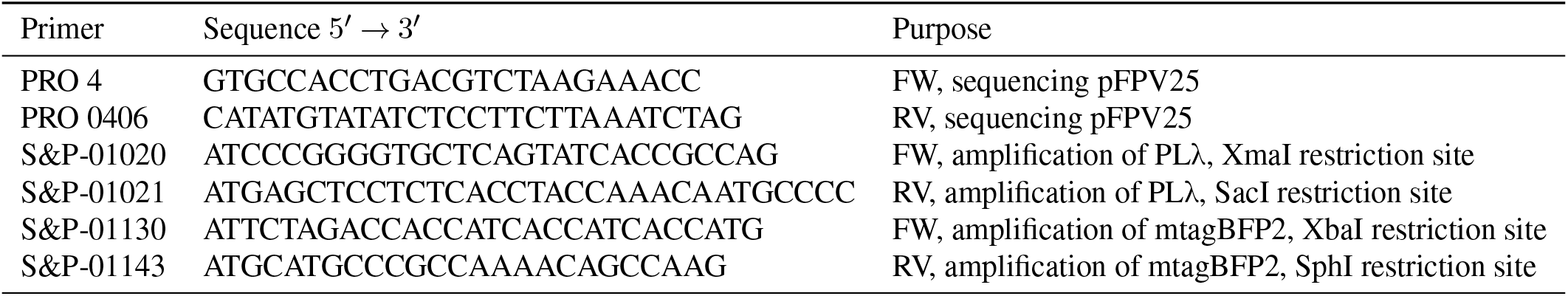
Oligonucleotides for plasmid construction (54).

### Biofilm assay and nanoparticle addition

The optical density of ONC of ATCC14028 mtagBFP2 was measured at 595 nm and corrected to *OD*_595_ = 2.5. These normalized cultures were further 10,000-fold diluted, corresponding to an initial bacterial density of approximately 2*e*5 cells/mL, in tryptic soy broth (TSB) diluted 5-fold for nutrient-rich conditions and 20-fold for nutrient-poor conditions. 396 µL of this suspension was added to µ-Slide 8 Well chambers (Ibidi) in addition to the appropriate concentration of ampiciline, and statically incubated for 48 h at 25 °C. For the visualization and measurement of EPS, EbbaBiolight 680 (Ebba Biotech AB) was added at the start of incubation using a 1,000-fold dilution following the manufacturer instructions.

After 48h of biofilm incubation, green fluorescent aminated (radius = 50 nm) or green fluorescent carboxylated (radius = 60 nm) polystyrene nanoparticles (Spherotech) were gently pipetted directly below the liquid-air interface to avoid structural disturbance of the biofilms, to a total of 4 µL of nanoparticle stock solution and thus final concentration of 10^−5^ w/v%.

### Image acquisition

Prior to single particle tracking, biofilms were imaged using an inverted fluorescence microscope (Z1 observer, Zeiss) with an 63x oil immersion objective at 6 µm above the well surface using an excitation wavelength of 450 nm 20 min after addition of the nanoparticles, these were tracked using frequency of 10 Hz during 50 s using an excitation wavelength of 495 nm.

For localization of nanoparticles with respect to bacteria, biofilms were imaged 1 hr after nanoparticle addition, which was performed identically to the single particle tracking experiments. Z-stacks of bacteria, EPS and nanoparticles were acquired simultaneously by respective excitation at 405 nm, 540 nm and 488 nm using a 63x oil immersion objective mounted on a confocal laser scanning microscope (LSM880, Zeiss). Z-stacks were captured on an Airyscan detector (Zeiss) using Fast Airyscan mode.

### Image processing

For the single particle tracking, blob detection was performed on every frame of the nanoparticle channel using the Trackmate plugin (55) implemented in the ImageJ platform (56) using a Difference of Gaussian filter with an estimated blob size of 1.5 µm. Detected spots with a quality metric below 20 were omitted from further analysis. The Linear Assignment Problem (LAP) tracker of Trackmate was used to link spots in subsequent frames allowing a maximal linking distance of 3 µm without gap closing.

After acquisition, Z-stacks were post-processed using Airyscan post-processing of Zen Black (Zeiss) with automatic Wiener Filter strength parameter. Nanoparticle Z-stacks were segmented using the Trackmate plugin as well (55), with quality threshold set to 100, and radius to 0.75 µm for aminated and 0.5 µm for carboxylated nanoparticles. For segmentation of bacteria, the signal in the Z-stacks was magnified using the histogram matching algorithm implemented in ImageJ (56) to match the intensity histogram of each slice to that at the bottom of the biofilm. Bacterial positions and geometry were extracted by splitting the Z-stacks in a set of substacks using a 4-by-4 in the xy-plane with a 20% overlap in both x and direction. A Hession-based Frangi vesselness filter was used to enhance blob-like features in each of the image substack, which were subsequently classified in bacteria and background using an Otsu threshold scaled with a factor 0.07. Binarized image substacks were stitched back together, followed by watershed segmentation of individual bacteria using the ImageJ platform (57). The position, radii and directions were obtained by computation of the 3D moment matrix of each individual blob (58). The largest eigenvalue was used as the length of the bacteria, while the two smaller radii were averaged out keeping the ellipsoid volume constant. Objects with a length smaller than 0.5 µm or radius smaller than 0.25 µm were omitted from further analysis. Finally, lengths bigger than 3 µm were set to 3 µm and the maximum radius was set 1 µm the same way. Thickness of the EPS was measured manually perpendicular to the bacterial cell wall of 50 randomly chosen bacteria in the central slice of the Z-stack (56).

### Brownian dynamics model

There are two separate entities in our model. The first are the time invariant spherocylindrical bacteria, with state variables of length *L*_*c*_, radius *R*_*c*_, node positions ***x***_0_ and ***x***_1_, matrix characteristic length *σ*_*M*_ and matrix viscosity Δ*η*_*M*_. The second entity are the nanoparticles, with state variables radius *R*_*p*_, mass density *ρ*_*p*_ position ***x***_*i*_(*t*) and experienced viscosity *η*(***x***_*i*_(*t*)). The environment state variables are temperature *T*, bulk density *ρ*_0_, nanoparticle concentration *C*_*p*_ and bulk viscosity *η*_0_. Nanoparticles experience reflective boundary conditions when they move too far from the biofilm. There is no interaction between nanoparticles. We assume the characteristic time of diffusion as an order of magnitude smaller than the characteristic time of biofilm growth (59), therefore assume the biofilm as static during the diffusion simulation. Diffusion is simulated for a total of 10 minutes, the degree of convergence over time is shown in Supplementary Fig. 9 and 10. State variables and their scales are listed in Table 2.

**Table 2.** Time-independent state variables when modelling nanoparticle diffusion in a biofilm environment. When *σ* _*M*_ is varied, we keep a constant Δ*η*_*M*_ = 10 mPa s, When Δ*η* _*M*_ is varied, we keep a constant *σ* _*M*_ = 0.5 µm.

| Symbol | Value [Default] | Unit | Description |
| --- | --- | --- | --- |
| $R_p$ | 60 | nm | Nanoparticle radius |
| $\rho_p$ | 1060 | kg/m<sup>3</sup> | Nanoparticle mass density |
| $R_c$ | 0.5 | $\mu\text{m}$ | Spherocylinder radius |
| $L_c$ | 3 | $\mu\text{m}$ | Spherocylinder length |
| $\sigma_M$ | 0.05 - 2 [0.5] | $\mu\text{m}$ | Matrix characteristic length |
| $\eta_0$ | 1 | mPa.s | Bulk viscosity |
| $\Delta\eta_M$ | 0 - 100 [10] | mPa.s | Viscosity at bacteria interface, in addition to $\eta_0$ |
| $k_{cp}$ | 20 | pN/$\mu\text{m}$ | Harmonic potential stiffness |
| $T$ | 300 | K | Bulk temperature |
| $C_p$ | 10 | pM | Nanoparticle concentration |
| $\rho_0$ | 997 | kg/m<sup>3</sup> | Bulk mass density |
| $T_s$ | 10 | minutes | Total simulation time |

Extracellular polymeric substances (EPS) interact with nanoparticles in the biofilm and slow down diffusion. This leads to inhomogeneities of the viscosity in the medium and thus the spatially varying overdamped Langevin equation

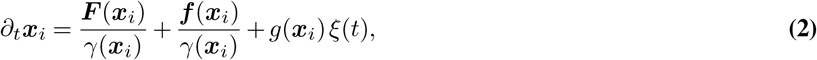

with *g*^2^(***x***_*i*_) = 2*k*_*B*_*T*/*γ*(*x*) the strength of Gaussian white noise *ξ*(*t*) with properties ⟨*ξ*(*t*)⟩ = 0 and ⟨*ξ*(*t;*)*ξ*(*t*^*′*^)⟩ = *δ*(*t− t*^*′*^), ***F*** (***x***_*i*_) the resultant of contact forces, *γ*(***x***_*i*_) the friction coefficient, *k*_*B*_ the Boltzmann constant, *T* the temperature and a drift force ***f***. The drift force originates from the Stratonovich convention, which according to Jacka and Oksendal best describes the diffusion of particles inside an inhomogeneous environment (60). Since nanoparticles are represented as spheres, we describe the friction coefficient with Stokes-Einstein so that *γ*(***x***_*i*_) = 6*πr*_*t*_*η*(***x***_*i*_), with *r*_*t*_ radius of the nanoparticle and *η*(***x***_*i*_) the local dynamic viscosity.

Since Gaussian viscosity kernels are often used for diffusion in heterogeneous environments (37), we will also assume that *η*(***x***_*i*_) declines according to a Gaussian with respect to the distance from the surface of the bacteria so that

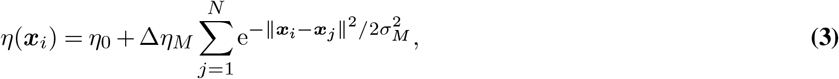

where Δ*η*_*M*_ is the difference between viscosity in water, *η*_0_ and viscosity near the surface of bacteria, ***x***_*j*_ the point on the surface of bacteria *j*, closest to the NP and *σ*_*M*_ the characteristic length scale of the viscosity kernel. The drift force 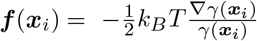 in Eq. 2 is then

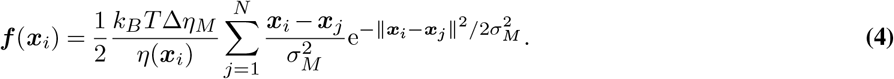

In addition, nanoparticles experience a gravity force *F*_*g*_ = *g*· (*ρ*_*p*_*− ρ*_0_) *V*_*p*_ towards the bottom of the biofilm, with *V*_*p*_ the volume of the spherical nanoparticle. Then, contact forces ***F*** (***x***_*i*_) between bacteria and nanoparticles are calculated as harmonic repulsive potentials, with stifness *k*_*cp*_. After contacts, experienced viscosity of each nanoparticle is calculated as described in Eq. 3. Resulting velocities and forces are calculated with the conjugate gradient method, after which resulting positions are calculated via a Forward-Euler integration scheme. Particles experience a closed boundary box surrounding the biofilm.

### Generation of virtual biofilms

We simulate biofilm structures using the Cahn-Hilliard equations

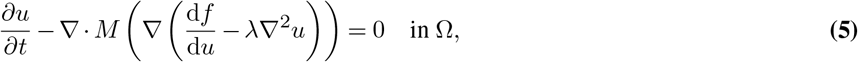

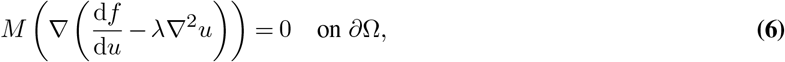

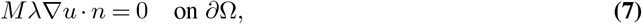

with the Dolfin platform from FEniCSx (61). We initialize the field of *u*(***x***) as

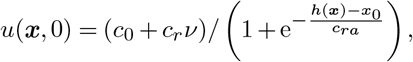

where *ν* is a uniformly distributed random number between -1 and 1, ***x*** is the voxel position in a 56 × 56 × 56 grid, *c*_*r*_ = 0.1, *c*_0_ = 0.5, *h*(***x***) is height at location ***x***, *x*_0_ = 0.15 and *c*_*ra*_ = 0.15. We simulate for 1*e*9 time steps with Dirichlet boundary conditions. The Dirichlet boundary conditions lead to lower mean of field *u*(***x***) over time, thus we multiply each *u*(***x***, *t*) value with ⟨*u*(***x***, *t*_*f*_)⟩ / ⟨*u*(***x***, *t*)⟩, with *t*_*f*_ the final time step, where ⟨*u*(***x***, *t*_*f*_)⟩ = 0.36. Since the characteristic length *L*_*t*_ scales with *t*^1/3^ (Lifshitz–Slyozov law), we generate biofilms at time steps 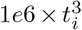, with *t*_*i*_ an integer from 0 to 10, such that 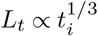. We seed bacteria at a constant volume density (2.8% v/v%), after which bacteria are accepted with probability *u*(***x***, *t*_*i*_), such that the final volume density is 1% v/v%. Characteristic length *L*_*t*_ of the virtual biofilms was calculated using Paraview contour filter, followed by the integrate variable filter to calculate surface area of the Cahn-Hilliard domains (62). Characteristic length was then calculated as *L*_*t*_ = *V*/*S*, with *S* the estimated surface and *V* the volume of the Cahn-Hilliard domain. Finally, the slope of *L*_*t*_ as a function of *t*^1*/*3^ was calculated with linear regression (following the Lifshitz–Slyozov law), such that *L*_*t*_ = 0 µm at *t*_*i*_ = 0, see Supplementary Fig. 11.

### Reaction-diffusion model in gyroid structures

The setup and reasoning for our finite element simulations of antibiotics diffusion in gyroid structure is explained in more detail in Supplementary information.

### Diffusion measures

The time averaged mean squared displacement (TAMSD)

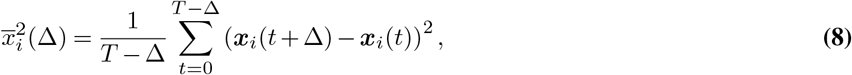

for particle *i*, where Δ is the lag time and *T* total track length. The diffusion coefficient *D* is calculated from the TAMSD via linear least squares, as 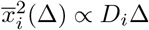. Diffusion exponent *α* is calculated from the TAMSD via nonlinear least squares, as 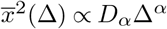.

The “affinity” and “coverage length” measures are computed by calculating pairwise surface-to-surface distances from each NP to each bacteria. The percentage of NPs which are closer than 1 µm to the closest bacteria are is called the affinity, while the median distance from each bacteria to the closest NP is called “coverage length”. A small coverage length indicates that most bacteria are well reached by the NPs and are likely susceptible to potential encapsulated treatments. It should be noted that as a treatment measure, the coverage length is expected to be dependent on both structure of the biofilm as well as on the concentration of NPs.

## Data availability statement

Data generated and analysed in this study are available on https://doi.org/10.48804/BTMCFO and source code for replication on https://gitlab.kuleuven.be/mebios-particulate/mpacts_biofilm_brownian_dynamics. Raw images are available upon reasonable request.

## Acknowledgements

We would like to thank Pauline Brepoels and Kenny Appermans for providing us with the *Salmonella enterica* Typhimurium strain and Bram Lories for insightful discussions. This work was supported by the FWO Flanders under grant agreements G046318N and the KU Leuven under grant agreements Research Fund C3/20/081, C24/18/046, RF 21/6343 SalmiPIG, PDM/20/123.

## References

1. Flemming, H.-C., Neu, T. R. & Wozniak, D. J. The EPS Matrix: The “House of Biofilm Cells”. Journal of Bacteriology 189:7945–7947 (2007).

2. Flemming, H. C. & Wuertz, S. Bacteria and archaea on Earth and their abundance in biofilms. Nature Reviews Microbiology 17:247–260 (2019).

3. Xiu, W. et al. Recent development of nanomedicine for the treatment of bacterial biofilm infections. View 2:20200065 (2021).

4. Fulaz, S., Vitale, S., Quinn, L. & Casey, E. Nanoparticle–Biofilm Interactions: The Role of the EPS Matrix. Trends in Microbiology 27:915–926 (2019).

5. Koo, H., Allan, R. N., Howlin, R. P., Stoodley, P. & Hall-Stoodley, L. Targeting microbial biofilms: Current and prospective therapeutic strategies. Nature Reviews Microbiology 15:740–755 (2017).

6. Karygianni, L., Ren, Z., Koo, H. & Thurnheer, T. Biofilm Matrixome: Extracellular Components in Structured Microbial Communities. Trends in Microbiology 28:668–681 (2020).

7. Peulen, T.-O. & Wilkinson, K. J. Diffusion of nanoparticles in a biofilm. Environmental science & technology 45:3367–3373 (2011).

8. Ikuma, K., Decho, A. W. & Lau, B. L. T. When nanoparticles meet biofilms—interactions guiding the environmental fate and accumulation of nanoparticles. Frontiers in microbiology 6 (2015).

9. Eleraky, N. E., Allam, A., Hassan, S. B. & Omar, M. M. Nanomedicine fight against antibacterial resistance: An overview of the recent pharmaceutical innovations. Pharmaceutics 12:1–51 (2020).

10. Gao, Y. et al. Size and Charge Adaptive Clustered Nanoparticles Targeting the Biofilm Microenvironment for Chronic Lung Infection Management. ACS Nano 14:5686–5699 (2020).

11. Leal, J., Smyth, H. D. & Ghosh, D. Physicochemical properties of mucus and their impact on transmucosal drug delivery. International Journal of Pharmaceutics 532:555–572 (2017).

12. Forier, K. et al. Transport of nanoparticles in cystic fibrosis sputum and bacterial biofilms by single-particle tracking microscopy. Nanomedicine 8:935–949 (2013).

13. Golmohamadi, M., Clark, R. J., Veinot, J. G. C. & Wilkinson, K. J. The role of charge on the diffusion of solutes and nanoparticles (silicon nanocrystals, nTiO2, nAu) in a biofilm. Environmental Chemistry (2013).

14. Sankaran, J. et al. Single microcolony diffusion analysis in Pseudomonas aeruginosa biofilms. npj Biofilms and Microbiomes 5:35 (2019).

15. Devlin, H. et al. A high throughput method to investigate nanoparticle entrapment efficiencies in biofilms. Colloids and Surfaces B: Biointerfaces 193:111123 (2020).

16. Witten, J. & Ribbeck, K. The particle in the spider’s web: transport through biological hydrogels. Nanoscale 9:8080–8095 (2017).

17. Li, X. et al. Control of nanoparticle penetration into biofilms through surface design. Chemical Communications 51:282–285 (2015).

18. Dunsing, V., Irmscher, T., Barbirz, S. & Chiantia, S. Purely Polysaccharide-Based Biofilm Matrix Provides Size-Selective Diffusion Barriers for Nanoparticles and Bacteriophages. Biomacromolecules 20:3842–3854 (2019).

19. Powell, L. C. et al. Quantifying the effects of antibiotic treatment on the extracellular polymer network of antimicrobial resistant and sensitive biofilms using multiple particle tracking. npj Biofilms and Microbiomes 7:13 (2021).

20. Chew, S. C., Rice, S. A., Kjelleberg, S. & Yang, L. <em>In Situ</em> Mapping of the Mechanical Properties of Biofilms by Particle-tracking Microrheology. Journal of Visualized Experiments 2015:1–8 (2015).

21. Schuster, B. S., Ensign, L. M., Allan, D. B., Suk, J. S. & Hanes, J. Particle tracking in drug and gene delivery research: State-of-the-art applications and methods. Advanced Drug Delivery Reviews 91:70–91 (2015).

22. Wang, B., Anthony, S. M., Bae, S. C. & Granick, S. Anomalous yet Brownian. Proceedings of the National Academy of Sciences 106:15160–15164 (2009).

23. Flemming, H.-C. & Wingender, J. The biofilm matrix. Nature Reviews Microbiology 8:623–633 (2010).

24. Lawrence, J. R., Swerhone, G. D. W., Kuhlicke, U. & Neu, T. R. In situ evidence for metabolic and chemical microdomains in the structured polymer matrix of bacterial microcolonies. FEMS Microbiology Ecology 92:fiw183 (2016).

25. Rodríguez-Suárez, J. M., Butler, C. S., Gershenson, A. & Lau, B. L. T. Heterogeneous Diffusion of Polystyrene Nanoparticles through an Alginate Matrix: The Role of Cross-linking and Particle Size. Environmental Science & Technology 54:5159–5166 (2020).

26. Powell, L. C. et al. Targeted disruption of the extracellular polymeric network of Pseudomonas aeruginosa biofilms by alginate oligosaccharides. npj Biofilms and Microbiomes 4 (2018).

27. Teirlinck, E. et al. Exploring Light-Sensitive Nanocarriers for Simultaneous Triggered Antibiotic Release and Disruption of Biofilms Upon Generation of Laser-Induced Vapor Nanobubbles. Pharmaceutics 11:201 (2019).

28. Stewart, P. S. & Franklin, M. J. Physiological heterogeneity in biofilms. Nature Reviews Microbiology 6:199–210 (2008).

29. Kosztolowicz, T., Metzler, R., Wasik, S. & Arabski, M. Modelling experimentally measured of ciprofloxacin antibiotic diffusion in Pseudomonas aeruginosa biofilm formed in artificial sputum medium. PLOS ONE 15:e0243003 (2020).

30. Stewart, P. S. et al. Conceptual Model of Biofilm Antibiotic Tolerance That Integrates Phenomena of Diffusion, Metabolism, Gene Expression, and Physiology. Journal of Bacteriology 201:1–24 (2019).

31. Wang, L., Hu, C. & Shao, L. The antimicrobial activity of nanoparticles: Present situation and prospects for the future. International Journal of Nanomedicine 12:1227–1249 (2017).

32. Hansing, J. et al. Nanoparticle filtering in charged hydrogels: Effects of particle size, charge asymmetry and salt concentration. The European Physical Journal E 39:53 (2016).

33. Blanco, E., Shen, H. & Ferrari, M. Principles of nanoparticle design for overcoming biological barriers to drug delivery. Nature Biotechnology 33:941–951 (2015).

34. Lin, C. C., Parrish, E. & Composto, R. J. Macromolecule and Particle Dynamics in Confined Media. Macromolecules 49:5755–5772 (2016).

35. Kondrat, S., Zimmermann, O., Wiechert, W. & Lieres, E. V. The effect of composition on diffusion of macromolecules in a crowded environment. Physical Biology 12:046003 (2015).

36. Stylianopoulos, T., Diop-Frimpong, B., Munn, L. L. & Jain, R. K. Diffusion anisotropy in collagen gels and tumors: The effect of fiber network orientation. Biophysical Journal 99:3119–3128 (2010).

37. Spakowitz, A. J. Transient Anomalous Diffusion in a Heterogeneous Environment. Frontiers in Physics 7:31–33 (2019).

38. Hartmann, R. et al. Emergence of three-dimensional order and structure in growing biofilms. Nature Physics 15:251–256 (2019).

39. Thomen, P., Valentin, J. D. P., Bitbol, A.-F. & Henry, N. Spatiotemporal pattern formation in E. coli biofilms explained by a simple physical energy balance. Soft Matter 16:494–504 (2020).

40. McNally, L. et al. Killing by Type VI secretion drives genetic phase separation and correlates with increased cooperation. Nature Communications 8 (2017).

41. Chubynsky, M. V. & Slater, G. W. Diffusing Diffusivity: A Model for Anomalous, yet Brownian, Diffusion. Physical Review Letters 113:098302 (2014).

42. Zhao, J. & Wang, Q. Three-Dimensional Numerical Simulations of Biofilm Dynamics with Quorum Sensing in a Flow Cell. Bulletin of Mathematical Biology 79:884–919 (2017).

43. Forier, K. et al. Probing the size limit for nanomedicine penetration into Burkholderia multivorans and Pseudomonas aeruginosa biofilms. Journal of Controlled Release 195:21–28 (2014).

44. Hansing, J. & Netz, R. R. Particle Trapping Mechanisms Are Different in Spatially Ordered and Disordered Interacting Gels. Biophysical Journal 114:2653–2664 (2018).

45. Zhou, H. & Chen, S. B. Brownian dynamics simulation of tracer diffusion in a cross-linked network. Physical Review E 79:021801 (2009).

46. Stewart, P. S. Theoretical aspects of antibiotic diffusion into microbial biofilms. Antimicrobial Agents and Chemotherapy 40:2517–2522 (1996).

47. Feng, J., Zhang, Z., Wen, X., Xue, J. & He, Y. Single Nanoparticle Tracking Reveals Efficient Long-Distance Undercurrent Transport in Upper Fluid of Bacterial Swarms. iScience 22:123–132 (2019).

48. Beroz, F. et al. Verticalization of bacterial biofilms. Nature Physics 14:954–960 (2018).

49. Simmons, E. L. et al. Biofilm Structure Promotes Coexistence of Phage-Resistant and Phage-Susceptible Bacteria. mSystems 5:1–17 (2020).

50. Deschênes, L. & Ells, T. Bacteria-nanoparticle interactions in the context of nanofouling. Advances in Colloid and Interface Science 277:102106 (2020).

51. Díaz-Pascual, F. et al. Breakdown of Vibrio cholerae biofilm architecture induced by antibiotics disrupts community barrier function. Nature Microbiology 4:2136–2145 (2019).

52. Cronenberg, T., Hennes, M., Wielert, I. & Maier, B. Antibiotics modulate attractive interactions in bacterial colonies affecting survivability under combined treatment. PLoS Pathogens 17:1–20 (2021).

53. Subach, O. M., Cranfill, P. J., Davidson, M. W. & Verkhusha, V. V. An enhanced monomeric blue fluorescent protein with the high chemical stability of the chromophore. PLoS ONE 6 (2011).

54. Valdivia, R. H. & Falkow, S. Bacterial genetics by flow cytometry: rapid isolation of Salmonella typhimurium acid-inducible promoters by differential fluorescence induction. Molecular Microbiology 22:367–378 (1996).

55. Tinevez, J. Y. et al. TrackMate: An open and extensible platform for single-particle tracking. Methods 115:80–90 (2017).

56. Schneider, C. A., Rasband, W. S. & Eliceiri, K. W. NIH Image to ImageJ: 25 years of image analysis. Nature Methods 9:671–675 (2012).

57. Preibisch, S., Saalfeld, S. & Tomancak, P. Globally optimal stitching of tiled 3D microscopic image acquisitions. Bioinformatics 25:1463–1465 (2009).

58. Ollion, J., Cochennec, J., Loll, F., Escudé, C. & Boudier, T. TANGO: A generic tool for high-throughput 3D image analysis for studying nuclear organization. Bioinformatics 29:1840–1841 (2013).

59. Picioreanu, C., Van Loosdrecht, M. & Heijnen, J. J. Effect of diffusive and convective substrate transport on biofilm structure formation: A two-dimensional modeling study. Biotechnology and Bioengineering 69:504–515 (2000).

60. Jacka, S. D. & Oksendal, B. Stochastic Differential Equations: An Introduction with Applications. Journal of the American Statistical Association 82:948 (1987).

61. Scroggs, M. W., Dokken, J. S., Richardson, C. N. & Wells, G. N. Construction of arbitrary order finite element degree-of-freedom maps on polygonal and polyhedral cell meshes. ACM Transactions on Mathematical Software 1:1–23 (2021).

62. Ayachit, U. The ParaView Guide: A Parallel Visualization Application, (Kitware, Inc., 2015).

