## Supplementary Information for "Spatial distribution of bacteria and extracellular polymeric substances impacts nanoparticle penetration in biofilms"

#### Supplemental Methods

**Reaction-diffusion model in gyroid structures.** To demonstrate the effect of macro-structural differences on the diffusion of antibiotics from NPs, we perform finite element method (FEM) diffusion simulations in an environment with high-absorption/low-diffusion regions, representing the EPS capturing antibiotics. With this setup, we want to illustrate how the spatial organization of EPS, for similar volume densities, impacts penetration, further supporting our statement that coarseness of diffusion-limiting structures can further limit penetration in the biofilm. As a reference distance, we use the diffusion length $L_d$. Parameters can be found in Table 3. We simulate diffusion within a gyroid structure, as it provides a simple mathematical representation of a porous structure in which the characteristic length can be tuned via a single parameter,

$$\mathcal{G}(\boldsymbol{x}) = \frac{2}{3}|\sin(x_a)\cos(y_a) + \sin(y_a)\cos(z_a) + \sin(z_a)\cos(x_a)| \equiv 0, \quad \textbf{(1)}$$

where $x_a = L_g x + \pi/16$, $y_a = L_g y + \pi/16$, $z_a = L_g z + \pi/16$ and $L_g$ a structural parameter determining the characteristic length scale of the gyroid. Note, that the scaling is chosen such that $\mathcal{G}(\boldsymbol{x}) \in [0,1]$ and the shift by the value $\pi/16$ was chosen such that the center of the antibiotics source is on a border of the gyroid surface. The antibiotics source acts as a point source $f(\boldsymbol{x})$, with a smooth transition from center to the edge, following a Gaussian decay as

$$f(\boldsymbol{x}) = \left(\frac{1}{25\pi L_d^2}\right)^{3/2} e^{-\left(\frac{x^2+y^2+z^2}{25L_d^2}\right)},$$

from the center of the domain. On the surface of the gyroid, diffusion will be slowest and capture rate highest, with a smooth transition between the interior surface and exterior bulk liquid. We define interior where $\mathcal{G}(\boldsymbol{x}) \leq v_f$. The transition from interior with low diffusion coefficient $D_e$ towards the exterior with high diffusion coefficient $D_aq$, is calculated as

$$D(\boldsymbol{x}) = D_{aq} + (D_e - D_{aq})\sqrt{\frac{\mathcal{G}(\boldsymbol{x}) - v_f}{1 - v_f}}.$$

The square root ensures a sharp transition from the center to the edge. The capture rate $r(\boldsymbol{x})$ transitions similarly as

$$r(\boldsymbol{x}) = r_{aq} + (r_e - r_{aq})\sqrt{\frac{\mathcal{G}(\boldsymbol{x}) - v_f}{1 - v_f}}.$$

Concentrations, absorption and diffusion rates are calculated with the DOLFIN Python package (1) in a cubic domain with dimensions $10\pi L_d$, discretized as a tetrahedral mesh ($99 \times 99 \times 99$). We solve the diffusion equation with source $f(\boldsymbol{x})$ and absorption $r(\boldsymbol{x})$

$$-\nabla\left[D(\boldsymbol{x}) \cdot \nabla u(\boldsymbol{x})\right] + r(\boldsymbol{x}) \cdot u(\boldsymbol{x}) = f(\boldsymbol{x}) \quad \textbf{(2)}$$

in steady state, as we are mainly interested in diffusion and absorption rates. Eq. 2 is solved with the generalized minimal residual method using incomplete LU factorization as preconditioner. $u(\boldsymbol{x})$ is the concentration at location $\boldsymbol{x}$, both sampled from a $P_1$ function space. The weak form of the equation Eq. (2) is then solved with natural boundary conditions at the edges of the box. A robustness check of $D_e$ and $\omega$ is shown in Supplementary Fig. 12.

#### Supplemental Tables

**Table 1.** Overview of biofilm density and height/thickness from CLSM images, with three biological repeats for each nanoparticle and nutrient availability combination.

| Nanoparticle | Nutrient availability | Volume density (v/v %) | Height (µm) |
|---|---|---|---|
| Aminated (Am) | Nutrient-poor | 3 ± 1 | 38 ± 4 |
| | Nutrient-rich | 7.2 ± 0.1 | 54 ± 8 |
| Carboxylated (Cb) | Nutrient-poor | 3.6 ± 0.6 | 32 ± 4 |
| | Nutrient-rich | 10 ± 4 | 62 ± 7 |

**Table 2.** Overview of multiple linear regression on coverage length in Fig. 4c.

| Variable | Estimate (µm) | Standard deviation (µm) | t-value | p-value |
|---|---|---|---|---|
| Intercept | 3.2 | 0.74 | 4.3 | $< 1e{-4}$ |
| $L_t$ | -0.012 | 0.037 | -0.32 | 0.75 |
| $\sigma_M$ | 50 | 1.4 | 34 | $< 1e{-4}$ |
| $L_t : \sigma_M$ | -0.89 | 0.072 | -12 | $< 1e{-4}$ |

**Table 3.** Parameters for gyroid finite-element diffusion simulations.

| **Name** | **Symbol** | **Value** | **Motivation** |
|---|---|---|---|
| Diffusion length | $L_d = 5\sqrt{\frac{D_e}{r_e}}$ | 1 | - |
| Gyroid length scale | $L_g$ | $[1-5]\,L_d$ | Sufficient difference between small and large $L_g$ |
| Void fraction | $v_f$ | 40% | Corresponds to free space in our experiments |
| Diffusion coefficient in low-diffusion regions | $D_e/D_{aq}$ | $10^{-2}$ | See Fig. 2c |

#### Supplemental Figures

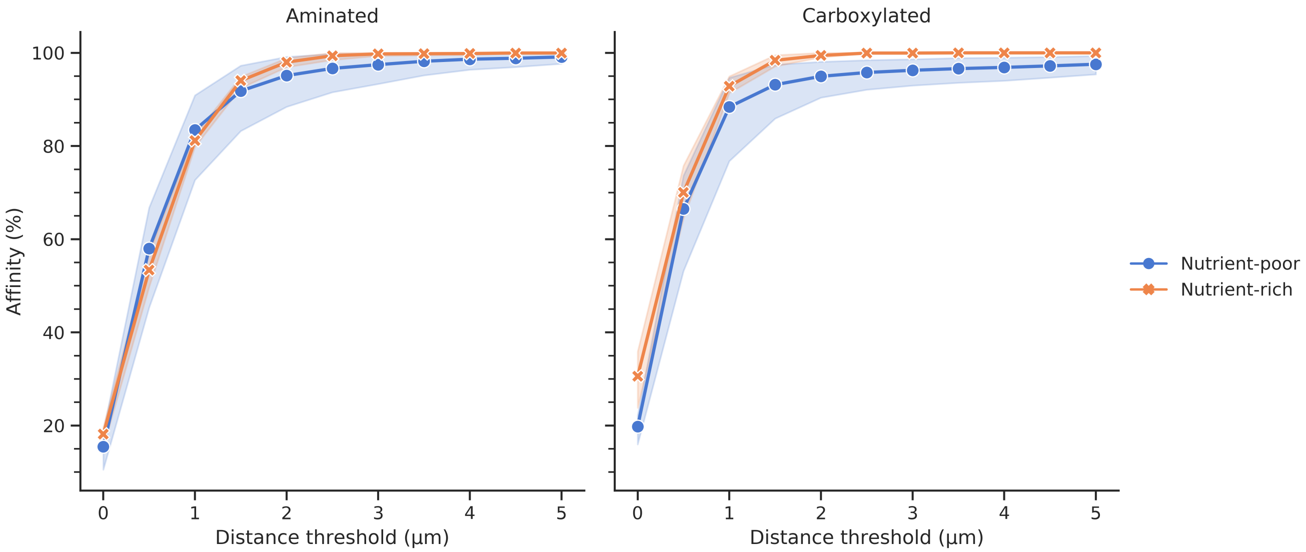


**Fig. 1.** Dependency of our affinity measure on the threshold value in *in vitro Salmonella enterica* Typhimurium biofilms.

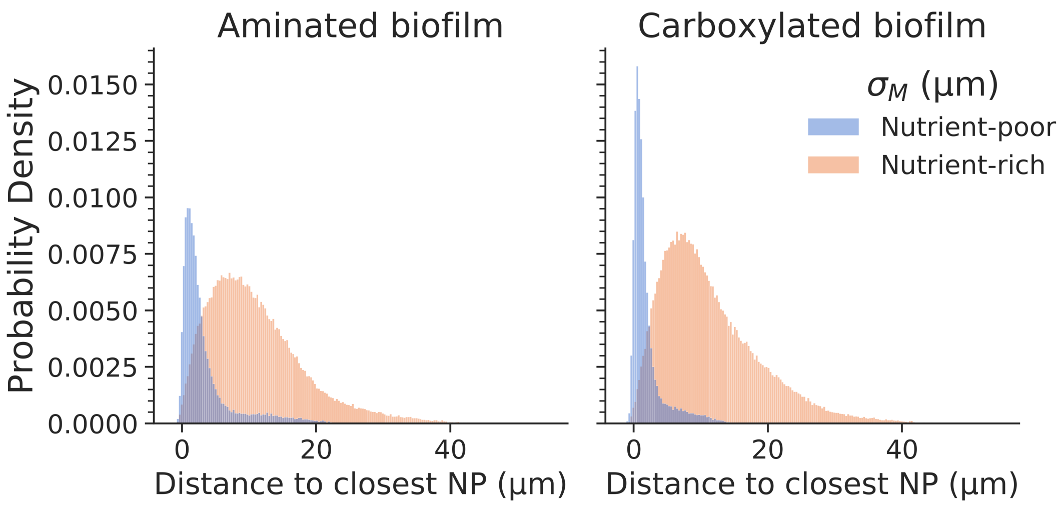


**Fig. 2.** Distributions for the distance of each bacteria to the closest NP. The median of these distributions is the coverage length. Data from CLSM experiments.

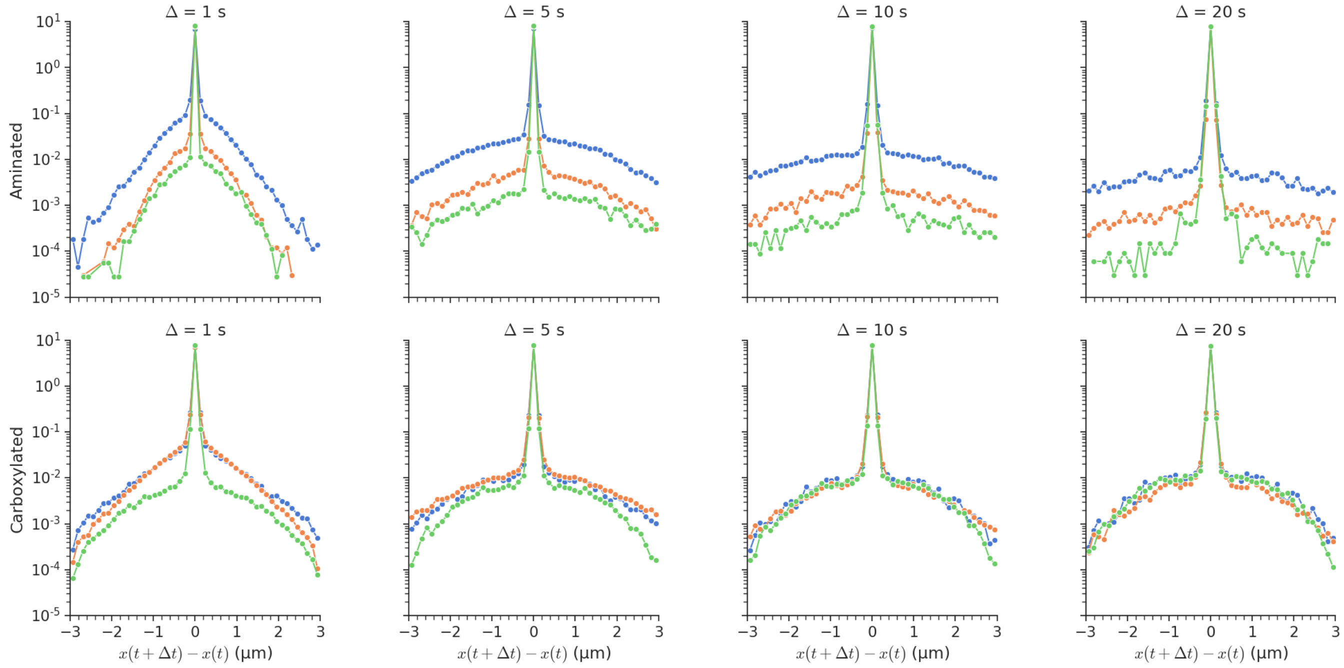


**Fig. 3.** Displacements curves for all SPT movies of both aminated and carboxylated nanoparticles, for various lag times $\Delta$. Different colors are different biological repeats.

### References


1. Logg, A. & Wells, G. N. DOLFIN: Automated finite element computing. *ACM Transactions on Mathematical Software* **37** (2010).

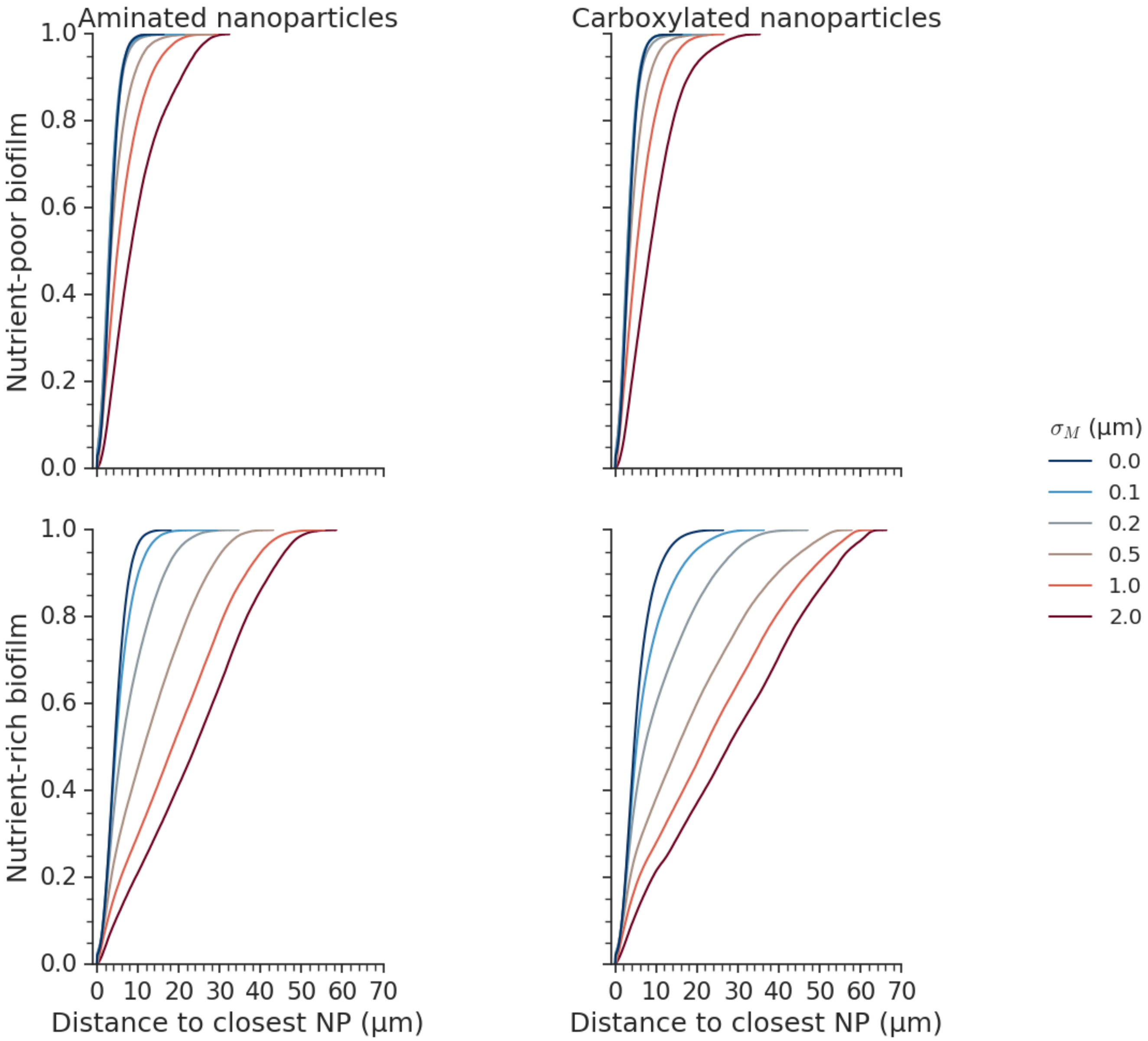


**Fig. 4.** Distributions for the distance of each bacteria to the closest NP. The median of these distributions is the coverage length. Data from simulations in segmented biofilms.

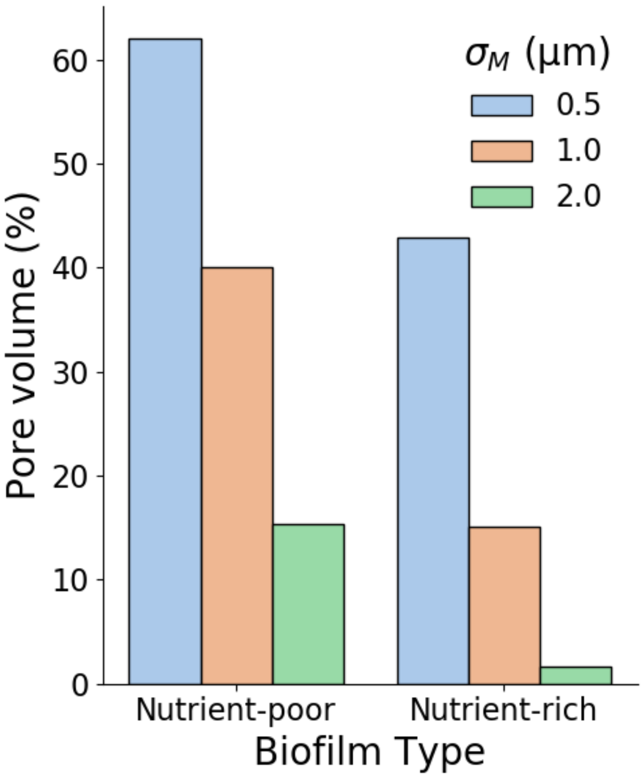


**Fig. 5.** Relative volume where $\eta(\boldsymbol{x})$ < 1.01 $\mathrm{mPa\,s} \approx \eta_0$ (porosity) of the biofilms shown in Fig. 3a and b.

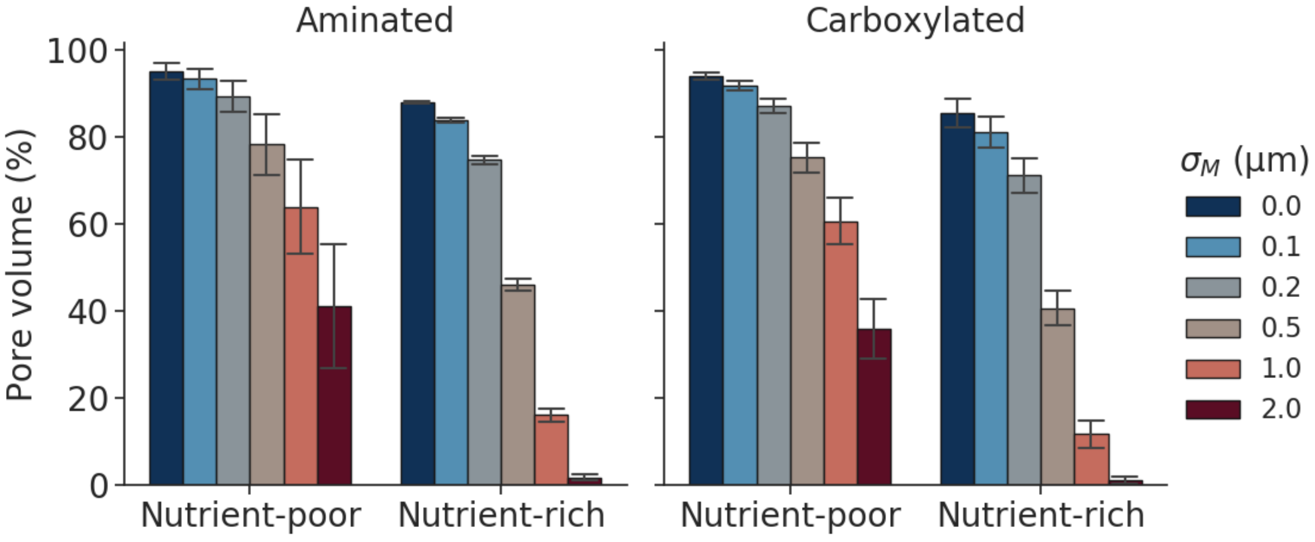


**Fig. 6.** Relative volume where $\eta(\boldsymbol{x})$ < 1.01 $\mathrm{mPa\,s} \approx \eta_0$ (porosity) of all imaged *in vitro Salmonella enterica* Typhimurium biofilms.

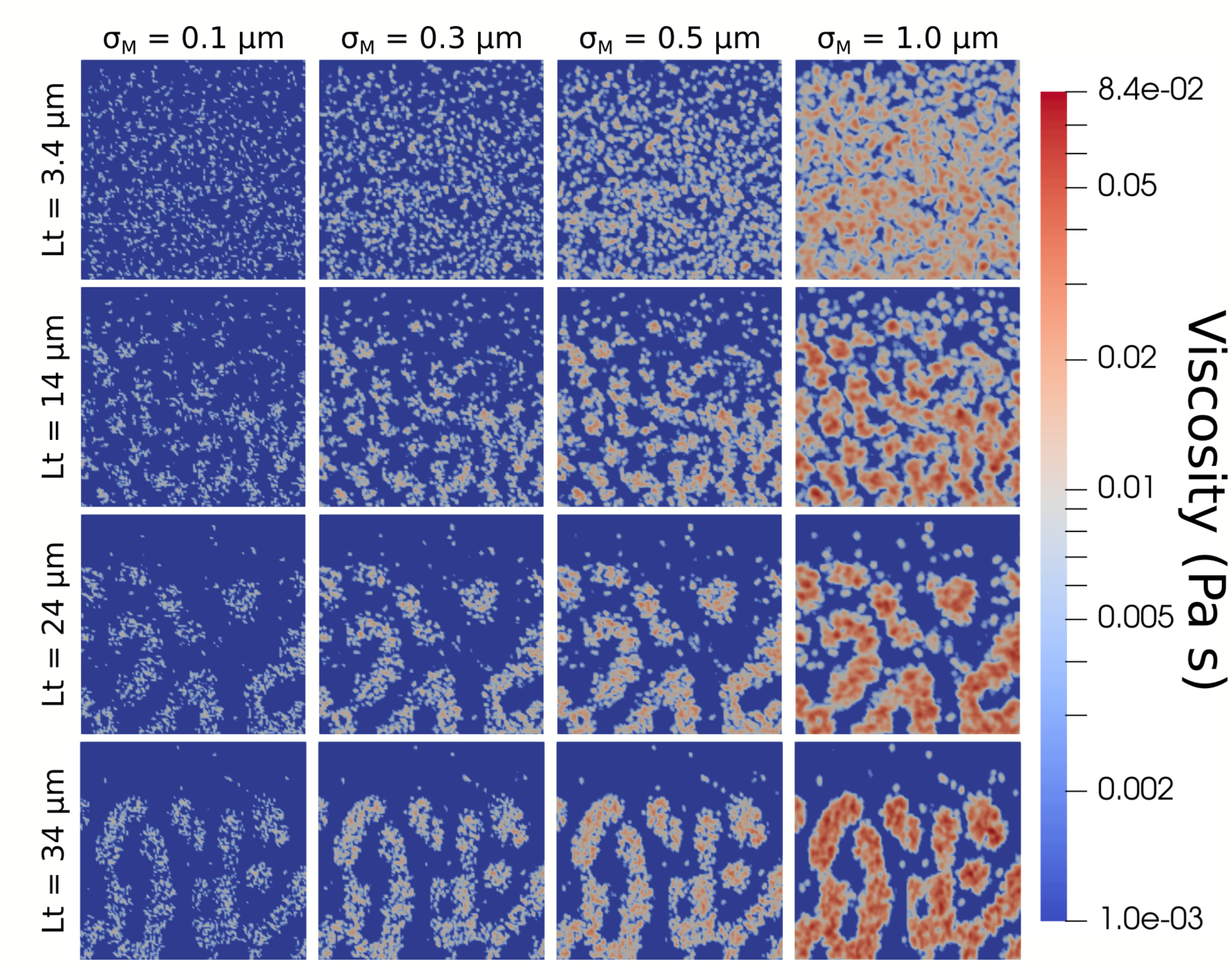


**Fig. 7.** XZ slice of viscosity in virtual biofilms of various characteristic length $L_t$ and matrix characteristic lengths $\sigma_M$. Slices are taken at the center of the biofilms.

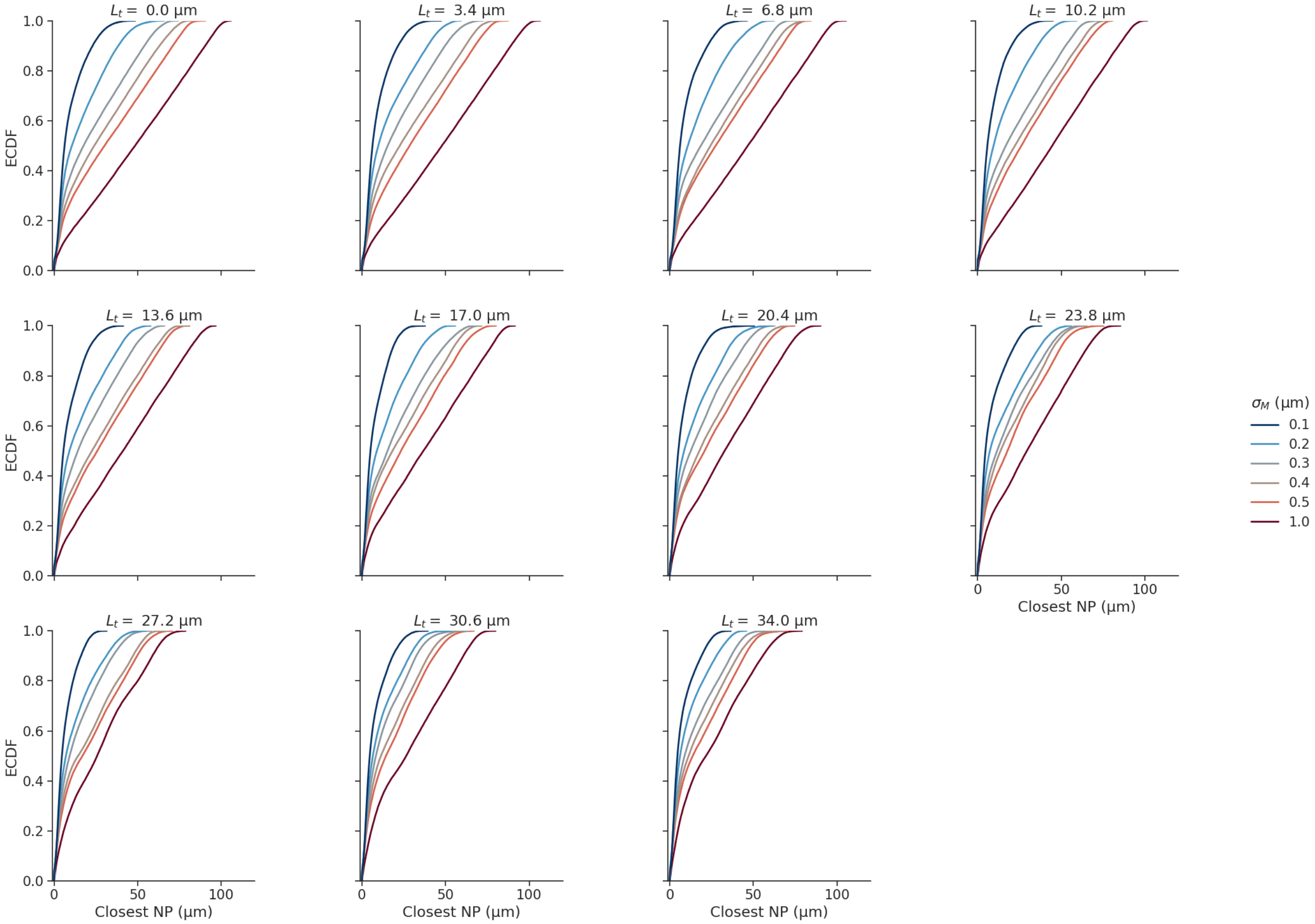


**Fig. 8.** Distributions for the distance of each bacteria to the closest NP. The median of these distributions is the coverage length. Data from simulations in virtual biofilms.

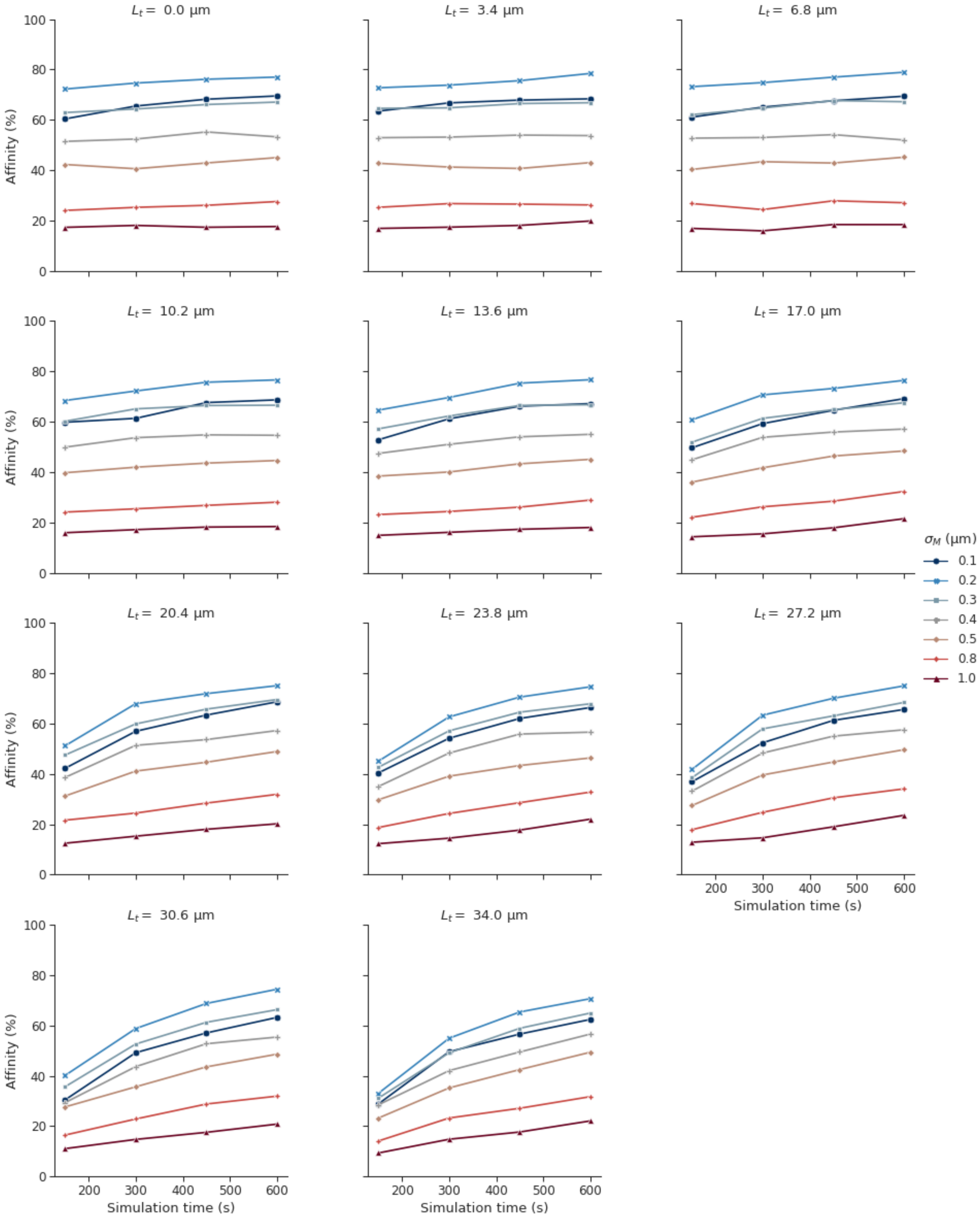


**Fig. 9.** Aging behaviour of affinity in virtual biofilm simulations, with a threshold of 0.3 $\mu$m.

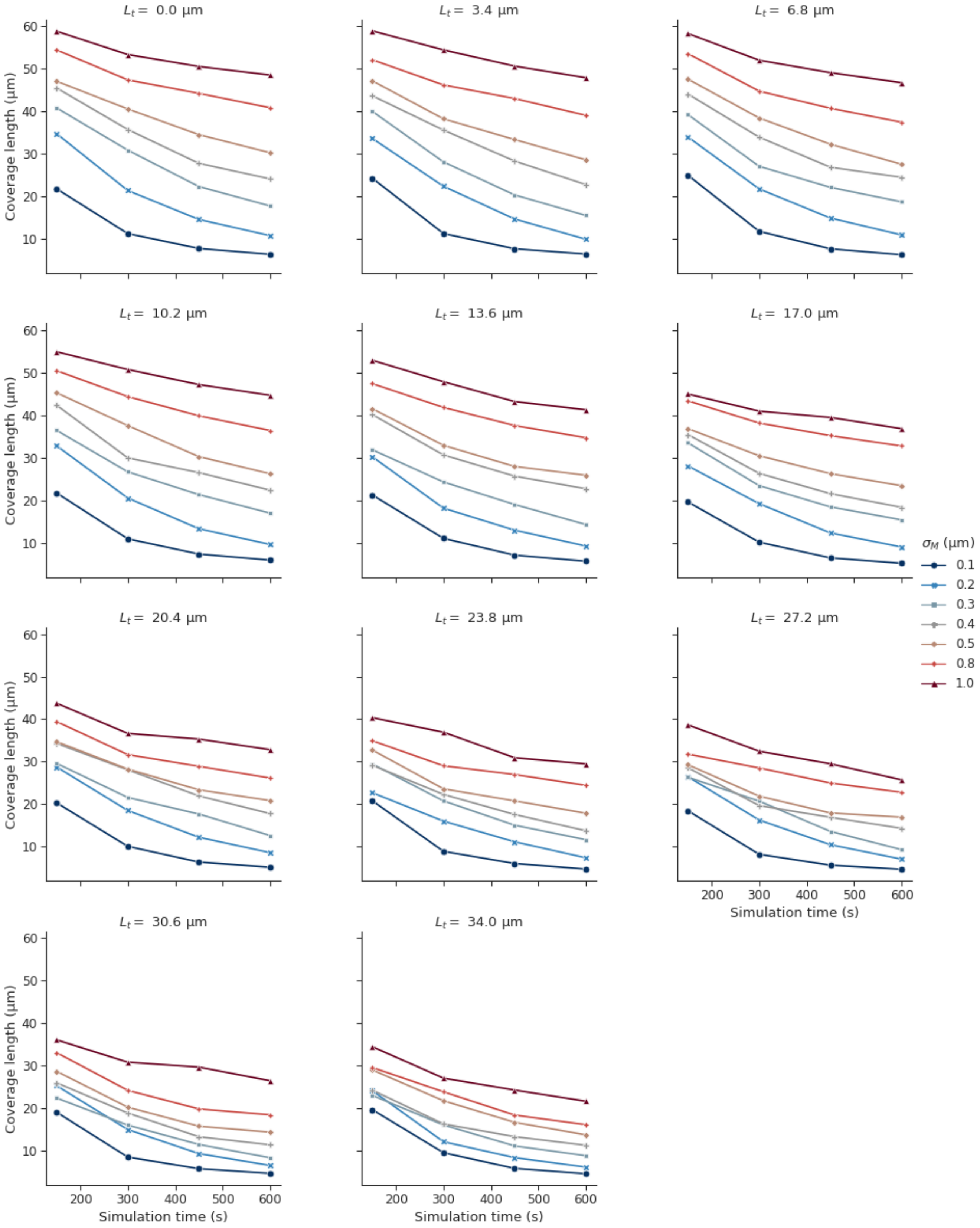


**Fig. 10.** Aging behaviour of the coverage length in virtual biofilm simulations.

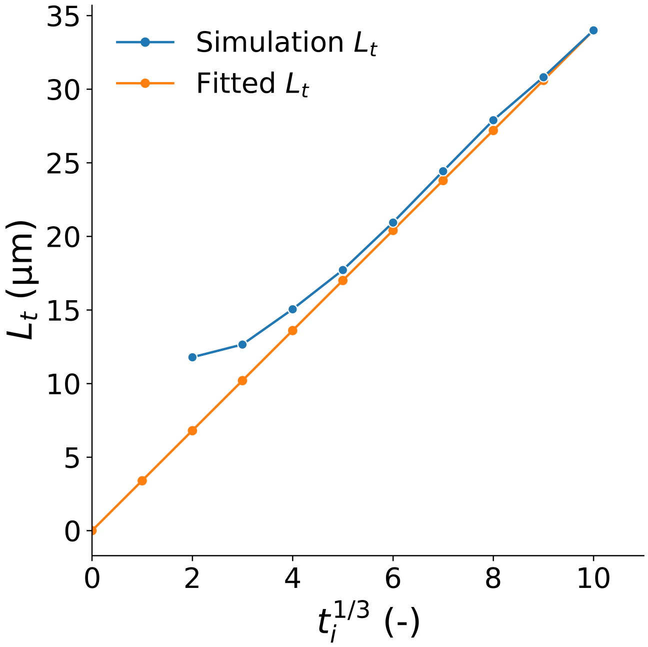


**Fig. 11.** Visualization of the calculation of characteristic length $L_t \propto t_i^{1/3}$, where $t_i$ is an integer from 0 to 10, such that time steps $1e6 \times t_i^3$ are taken from the Cahn-Hilliard simulations. At small $t_i$, $L_t$ diverges from the linear behaviour as the surface cannot be calculated accurately anymore for small characteristic lengths.

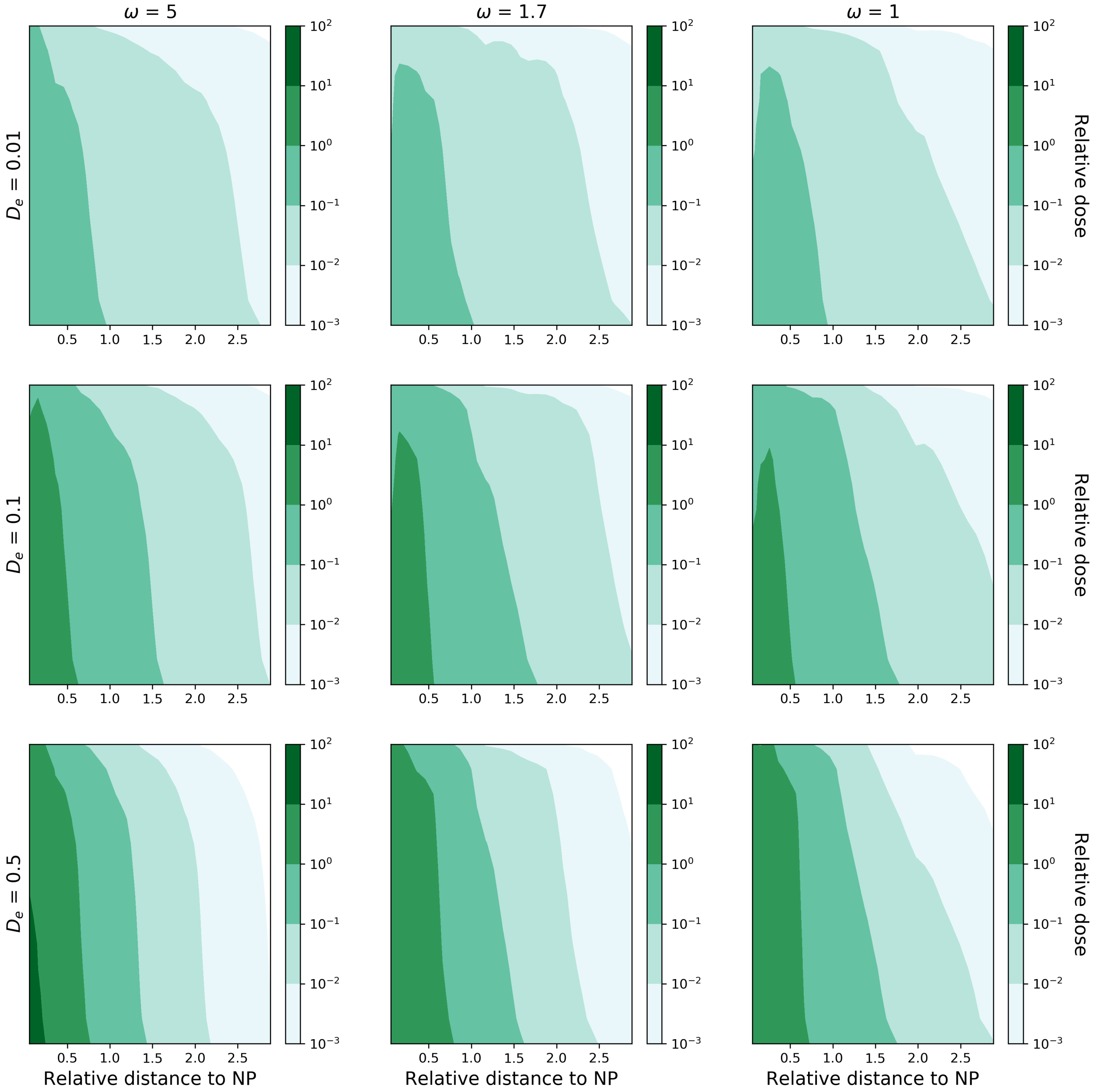


**Fig. 12.** Robustness study for the FEM diffusion simulations in a gyroid structure from Fig. 4e, with $v_f = 0.4$ and $r_e = 0.25$.
